# Temporal multi-omic profiling of immune, gut, and microbiome responses to ischemic stroke reveals convergence of host and microbial perturbations one week after brain injury

**DOI:** 10.64898/2026.05.25.727504

**Authors:** Junjie Guan, Burak Kizil, Garima Kalakoti, Delf-Magnus Kummerfeld, Oleksii Doroshenko, Vanesa Pelcastre-Neri, Nadine Chiara Frigger, Emilio Cirri, Nadine Pömpner, Manisha Goyal, Christina Janster, Johannes Zimmermann, Handan Melike Dönertaş, Katarzyna Winek

## Abstract

Ischemic stroke poses a significant medical challenge with limited therapy options, therefore detailed understanding of stroke-associated pathophysiological processes across systems and organs is crucial for further research and identification of novel therapeutic targets. In this manuscript, we provide parallel multi-omic host and gut microbiome characterization on several timepoints (day 1, 7 and 14) in the mouse experimental stroke model (middle cerebral artery occlusion, MCAo). Expanding existing host-derived datasets, we profiled transcriptomes from microglia, brain-infiltrating leukocytes and peripheral leukocytes using single cell RNA sequencing. Our data deliver time-resolved characterization of microglial subtypes and highlight heterogeneous dendritic cell populations as main interaction partners of microglia on all timepoints. In peripheral blood, we did not observe large transcriptomic differences when comparing the immune subsets from MCAo and sham-operated control animals. Here, the neutrophils exhibited most transcriptomic changes on day 1 among all blood leukocytes. Parallel proteomic analysis of 5 intestinal segments (duodenum, jejunum, ileum, caecum and colon) and mesenteric lymph nodes highlighted day 7 as the most important timepoint for changes in the gut-related metabolic pathways especially in the jejunum and colon. Specific hypothesis testing revealed compartmentalized regulation of gut-related immune pathways and proteins related to gut permeability. Finally, gut microbiome analyses (longitudinal metatranscriptomics including day -1, 3, 7, 14, and metagenomics from day 14) highlighted temporally matched changes in microbial gene expression (with day 7 emerging again as the most relevant timepoint), larger overall community perturbations when compared to baseline from day -1 in stroke animals and expansion of facultative anaerobes on day 7.

**Graphical abstract:** We performed longitudinal multi-omic host and gut microbiota profiling from experimental stroke and sham control mice. Our data indicate marked differences in brain-infiltrating leukocytes and microglia already on early timepoints after surgery, whereas stroke-specific changes in gut proteomics, bacterial gene expression and community structure emerge on day 7 after middle cerebral artery occlusion/sham surgery.

Created in https://BioRender.com

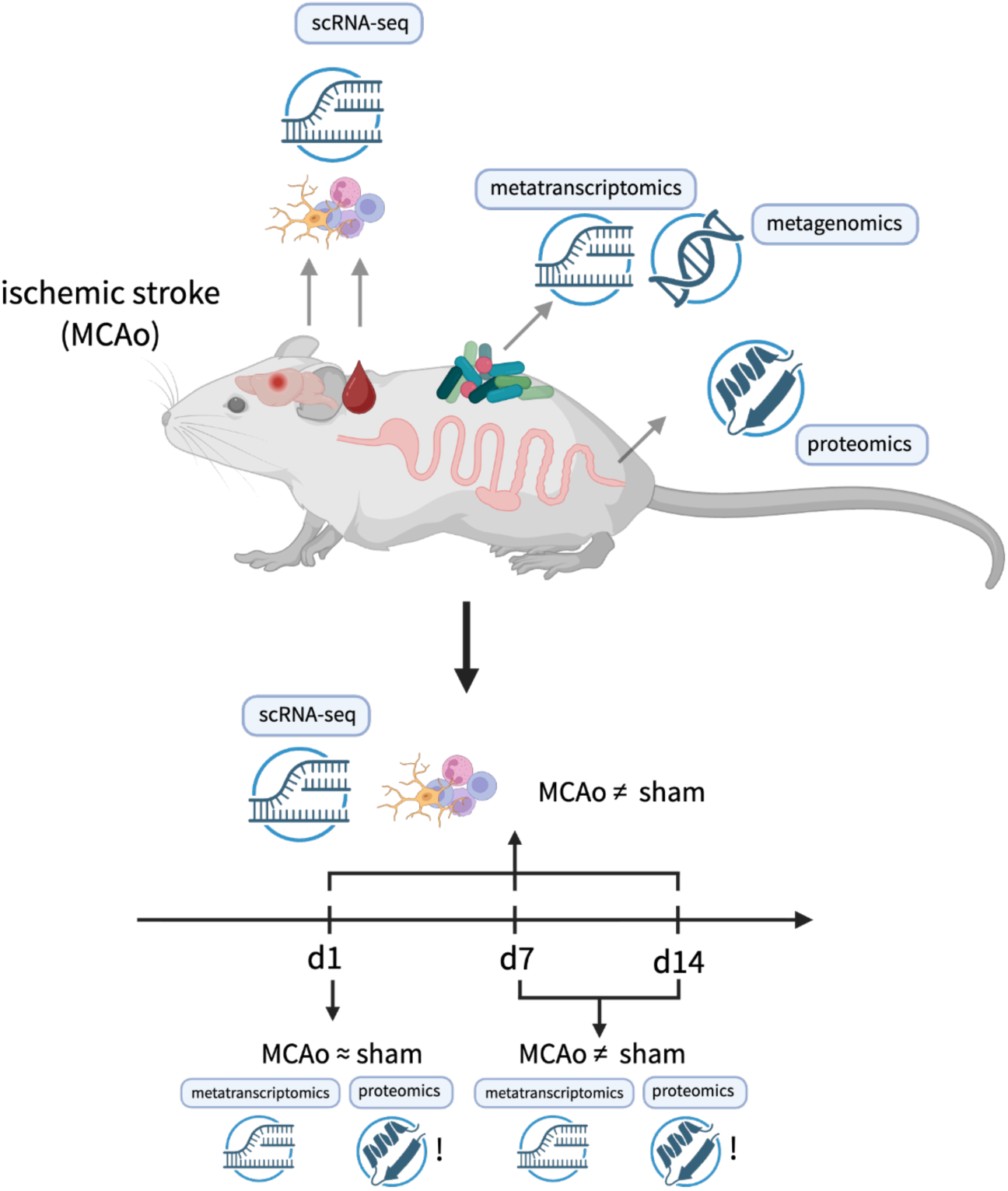

## Introduction

Despite significant progress in medical care and research, stroke is still one of the leading causes of death and disability worldwide ^1^. The immune system is one of the main contributors to stroke pathogenesis and a modifier of stroke outcome, as shown by numerous studies investigating the roles of diverse immune subpopulations in experimental models ^2–4^. Microglia, the brain-resident macrophages, respond first to the ischemic injury, transitioning from their homeostatic surveilling state to the activated form. Microglia migrate to the injury site and initiate the inflammatory response producing reactive oxygen species (ROS), cytokines/chemokines and metalloproteinases ^5^. Experimental studies show that depletion of microglia is generally detrimental in stroke models ^5^ as they participate in clearing up the debris, restrain astrocyte responses to the injury and produce factors boosting the repair processes. Neutrophils, the first peripheral immune cells to arrive at the injury site in the brain with numbers peaking in the first days after stroke, further drive inflammatory process and are followed by monocytes differentiating to macrophages and dendritic cells (DCs), natural killer (NK) cells and cells of the adaptive immunity - T cells and B cells ^6,7^. These findings are supported by clinical observations, although detailed knowledge about temporal dynamics and specific roles of immune subpopulations, especially in the clinical scenario, is limited. While the brain is a place of inflammatory response, stroke induces changes in peripheral organs and tissues. A prominent example is the suppression of immune functions in the periphery, which contributes to increased susceptibility to infections ^8,9^. Recent emergence of the novel research area focusing on reciprocal effects of stroke on the gut microbiota further highlights the importance of brain-body signaling after stroke ^10^. The gut microbiota, the largest collection of commensals in the body, has been identified as another factor impacting prognosis after stroke and contributing to disease susceptibility. Several studies reported rapid dysbiosis after cerebral ischemia ^11,12^ that can extend to later timepoints ^13^. Changes in gut motility and especially permeability and the question of potential translocation of bacteria or bacterial products after stroke remain controversial ^11,13–17^. Furthermore, recent studies provide early data on longitudinal profiling of the gut microbiota after brain injury ^13^ and gut microbiota emerges as an essential regulator of immune responses after ischemic stroke ^18^. However, microbial function (rather than composition alone) and its temporal coupling to host intestinal changes, particularly at the protein level, have not been characterized after stroke. In this manuscript, we aim to close this gap and provide a comprehensive characterization of the host (single cell transcriptomics from microglia, brain-infiltrated leukocytes and blood cells, proteomics from intestinal sections and mesenteric lymph nodes) and gut microbiota (metatranscriptomics and metagenomics) in response to ischemic stroke over several timepoints spanning the acute (d1) and subacute phases (d7 and d14) after brain injury. We identified rapid, stroke-specific changes in microglia and brain-infiltrating cells, while alterations in the intestinal compartments, microbial composition and function were mostly overshadowed by the effects of the surgical intervention on day 1. At this timepoint, changes in pathways related to T cell activation highlighted the ileum as a relevant immune compartment. Overall, however, day 7 emerged as a critical timepoint for compartment-specific host metabolic reorganization in the gut coupled with changes in the bacterial transcriptome and rearrangements in the bacterial community featuring expansion of facultative aerobes.

Our study delivers a unique combination of multiomic readouts from a single organism, provides the first post-stroke metatranscriptomic dataset and a solid framework to further dissect brain–immune–gut interactions after stroke.

## Results

To characterize host and gut microbiota responses to ischemic stroke, we employed mouse middle cerebral artery occlusion (MCAo), a widely used experimental stroke model. We conducted multi-omic profiling of host and gut microbiota responses to ischemic stroke over acute and subacute phase after injury. On the host side, to investigate the transcriptomic profiles of immune cells involved in post-stroke sequelae, we performed single-cell RNA sequencing (scRNA-seq) of FACS-sorted microglia and leukocytes isolated from the brain, as well as leukocytes from peripheral blood (Supplementary Fig. 1a-b). Cells were collected on day 1, 7 and 14 after MCAo induction. Simultaneously, gut sections and mesenteric lymph nodes (MLN) were collected for proteomic analyses on d1, d7 and d14 after MCAo. From a subcohort of animals (endpoint d14), stool pellets were collected one day before the surgery (d-1), d1, d3, d7 and d14 thereafter. Metagenomics was performed from fecal samples collected on d14 (see Figure 1a for the experimental design).

**Figure 1.**
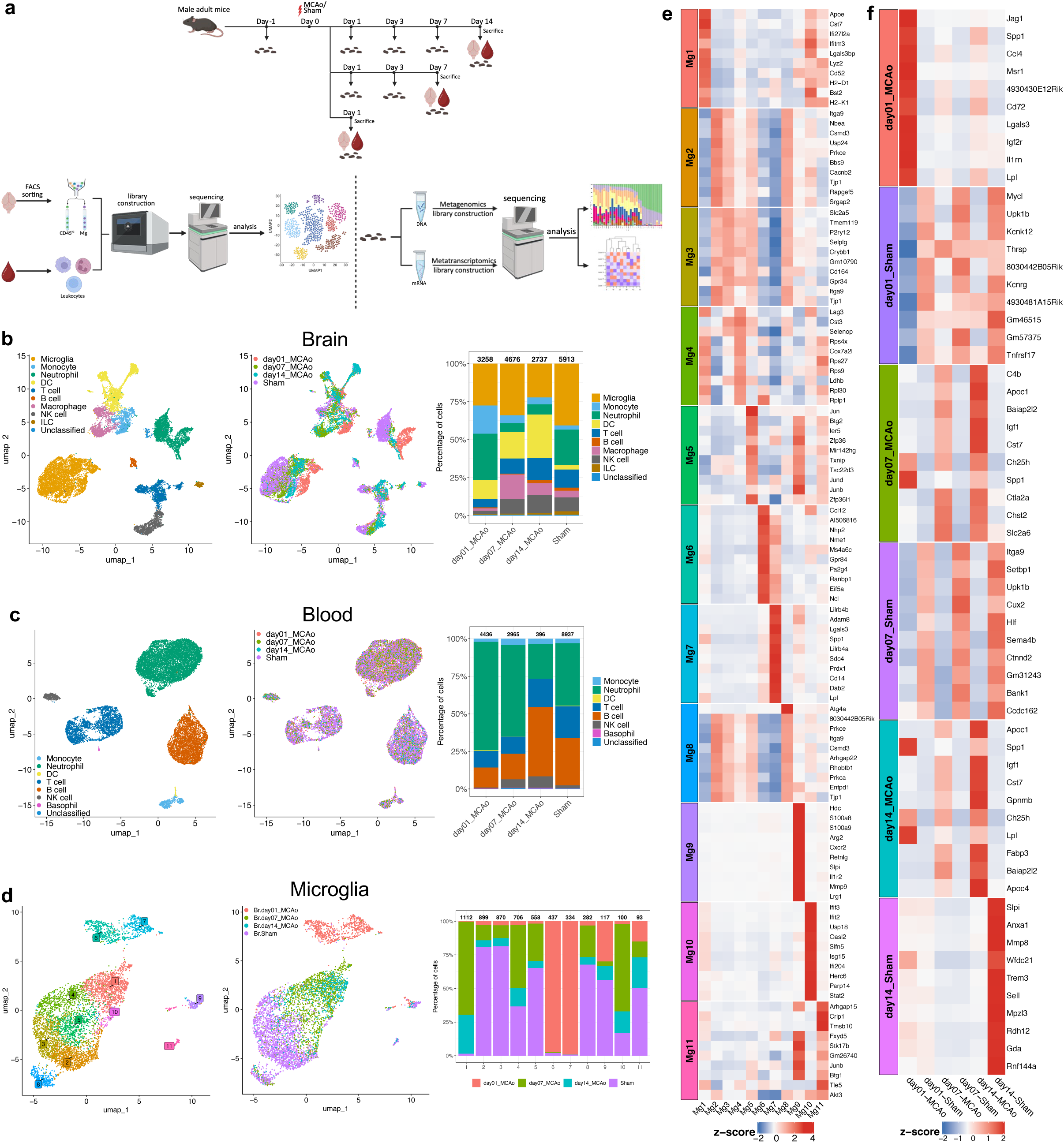
Microglia transcriptional programs shift from acute neuroinflammatory response to specialized immune programs. **(a)** Schematic overview of the experimental design and multi-omics workflow. Adult mice underwent middle cerebral artery occlusion (MCAo) or sham surgery on day 0. Fecal samples were collected longitudinally before surgery and at the indicated time points after surgery. At days 1, 7, and 14, brain and peripheral blood samples were harvested. Brain cells were dissociated to isolate leukocytes (CD45^hi^) and microglia (Mg) by flow cytometry sorting. Blood leukocytes were isolated after the lysis of red blood cells. Isolated cells were subjected to single-cell RNA sequencing (scRNA-seq). In parallel, fecal samples were processed for metagenomic and metatranscriptomic sequencing, followed by downstream bioinformatic analyses. **(b, c)** *Left*: Uniform manifold approximation and projection (UMAP) of brain (b) or blood (c) immune cells, colored by annotated major cell types; *Middle*: UMAP of the same brain (b) or blood (c) cells colored by experimental groups, including day01_MCAo, day07_MCAo, day14_MCAo, and pooled Sham; *Right*: Stacked bar plot showing the relative proportions of major immune cell types in brain (b) or blood (c) across experimental groups, numbers above the bars indicate the total number of cells in each group. **(d)** *left*: UMAP of microglia subset from brain transcriptomes reveals eleven clusters; *middle*: UMAP of the same microglia colored by experimental groups; *right*: Stacked bar plot showing the relative proportions of experimental groups across clusters, numbers above the bars indicate the total number of cells in each cluster. **(e)** Heatmap of top 10 differentially expressed genes (DEGs) across microglial clusters. For each microglial cluster (Mg1 – Mg11), DEGs were identified by comparing that cluster with all other microglial cells. Top 10 upregulated genes in each cluster are shown in the heatmap. **(f)** Heatmap of top 10 time point-specific microglial DEGs in MCAo versus sham mice. Differential expression analysis was performed separately at day 1, day 7, and day 14 by comparing microglia from MCAo and sham groups within each time point. Top 10 DEGs from each comparison are shown and grouped by the condition in which they were enriched. Heatmap rows represent genes and columns represent clusters (e) or experimental groups (f). Color intensity indicates scaled average expression (z-score). Top 10 DEGs were selected based on |log2FC| ≥ log2(1.5), pct ≥ 0.75 (e) or 0.1 (f), and Bonferroni-adjusted p < 0.05, ranked by log2FC. Panel (a) was created in https://BioRender.com

In the scRNA-seq analysis, after unsupervised clustering and uniform manifold approximation and projection (UMAP) 16,584 cells from the brain were identified and classified by unsupervised cell type annotation and expression of established marker genes as following 9 major cell types: microglia, monocytes, dendritic cells (DCs), macrophages, neutrophils, T cells, B cells, natural killer (NK) cells, innate lymphoid cells (ILCs), plus one unclassified cluster (Fig. 1b, Supplementary Table 1). Similarly, we identified 7 major cell types in 16,734 blood cells, including neutrophils, B cells, T cells, NK cells, monocytes, DCs, basophils and one unclassified cluster (Fig. 1c). We provide below detailed characterization of microglia and DCs as one of the main microglia interaction partners in the post-stroke brain, and finally peripheral blood neutrophils showing the most pronounced transcriptomic changes at the acute timepoint.

### Microglia shift from inflammatory activation to disease-associated and interferon-responsive states

Microglia act as the first responders to brain injury ^19^. To gain further insights into the effect of stroke on transcriptional programs in microglia, we have performed unsupervised clustering for microglial cells only, resulting in 11 clusters, displaying distinct sham and MCAo clusters (Fig. 1d). Clusters 2, 3 and 8 were predominantly derived from sham mice (with 18.6% to 32.2% of cells from MCAo animals) and expressed canonical homeostatic microglia markers (e.g., P2ry12, Tmem119, Gpr34), indicating presence of homeostatic microglia (Fig. 1e and STab. 2 for differentially expressed genes (DEGs) of each cluster). Cluster 2 and cluster 8 showed additional enrichment of adhesion- or interaction-associated transcripts (e.g., Itga9, Entpd1, and Tjp1), pointing to a contact-remodeling microglial state ^20,21^. Atg4a in cluster 8 is a key autophagy regulator, and microglial autophagy has been implicated in modulating neuroinflammatory responses after ischemic stroke, which suggests that enhanced autophagy-related transcription may influence post-ischemic immune remodeling ^22,23^. Day 1 post-stroke microglia dominated cluster 6 and cluster 7. Cluster 6 was characterized by genes related to chemokine and complement (e.g., Ccl12, Gpr84, and C5ar1), which highlights complement-microglia signaling as an early component of post-stroke neuroinflammation ^24,25^.

Other prominent genes in this cluster are related to upregulation of translational programs and proliferation (Ncl, Nhp2, Pa2g4, Eif5a, Ranbp1, Nme1) ^26^.

Cluster 7 exhibited an osteopontin-associated phagocytic program (e.g., Spp1, Lgals3, Lpl, and Cd14) and expression of antioxidant Prdx1 ^27^. These phagocytosing states resemble microglial populations reported in stroke models, consistent with early debris clearance and tissue remodeling after ischemic injury ^6,28–30^. Additionally, the enriched expression of Lilrb4a and Lilrb4b implies an engagement of neuroprotective transcriptional programs as Lilrb4b upregulation in microglia has been shown to limit brain-infiltration of damage-mediating CD8+ cells ^31^. A similar microglial cluster, expressing Lilrb4, Lgals3, Lpl and Spp1, has been identified by Ma et al. ^31^. Cluster 9 was not further annotated because its top markers were dominated by neutrophil-associated genes, suggesting possible contamination or doublets. Cluster 11 was not functionally interpreted because its marker profile was low-specificity and did not define a robust microglial activation state.

Day 7 and 14 stroke microglia were found mostly in cluster 1 and cluster 10. Cluster 1 showed high expression disease-associated microglia (DAM) genes including Apoe, Cst7, and Lyz2 ^6,30^ with simultaneous increased expression of MHC-I components (H2-D1, H2-K1, and B2m) and interferon-induced genes (Ifi27l2a, Ifitm3, Bst2) ^32,33^. Marked Ifi27l2a upregulation at 14 days post-stroke has been previously found in aged brains and Ifi27l2a identified as central regulator of the microglial pro-inflammatory responses ^32^. Our data strengthens this observation and extends it by describing cluster 1 as prominent on day 7 after stroke. Cluster 10 showed robust type I interferon-stimulated gene signature (e.g., Ifit2, Ifit3, Isg15, and Stat1), which corresponds to an interferon-responsive microglial (IRM) state. Type I interferon (IFN-I) signaling is activated following ischemic stroke, with microglia showing prominent interferon-stimulated gene expression that may perpetuate neuroinflammation during the subacute phase ^33–35^. Cluster 4 comprised mixed d7 and d14 post-stroke and sham microglia and exhibited enrichment of broadly expressed microglial transcripts (Cst3, Selenop) together with increased Lag3 expression. Given recent evidence linking Lag3 to regulation of microglial activation and DAM-associated programs, this cluster may represent microglia transitioning from homeostatic toward reactive states. ^36^. Finally, cluster 5, dominated by sham-microglia with contribution of cells from d7 and d14 after stroke showed expression of immediate-early genes (e.g., Jun, Junb, Btg2, Zfp36) which are involved in acute-response and stress-induced transcriptional activation ^37,38^. This phenotype could be related to our sample processing pipeline and FACS-sorting inducing expression of stress-associated transcripts ^38,39^.

Additional comparisons between stroke and sham microglia from respective timepoints (Fig. 1f with top 10 DEGs and whole list in STab. 3) revealed acute injury-associated transcriptional program characterized by strong chemokine induction (Ccl2, Ccl3, Ccl4, and Ccl12), inflammatory feedback control (Il1rn), scavenger/phagocytic and lipid-handling genes (Msr1, Spp1, Lpl, Lgals3, Dab2, Plin2, and Igf2r), and redox-stress genes (Prdx1, Glrx, Srxn1, and Mt2) ^19,40^ described previously as “stroke associated microglia” ^27^. By d7, MCAo microglia had shifted toward a phagocytic and lipid-remodeling phenotype enriched for Spp1, Apoe, Apoc1, Apoc2, Igf1, Lpl, Fabp5, Cst7, Clec7a and Ch25h, consistent with DAM-like repair-associated microglia described in recent stroke transcriptomic studies ^32,41–43^. This profile coexisted with a type I interferon module (lfi27l2a, Ifit1, Zbp1, Rsad2, Oasl2, Oas1a, and Oas2), indicating that reparative and interferon-responsive states are simultaneously present at this stage ^32,41,43,44^. At day 14, this injury-associated program remained prominent with a partial recovery of canonical homeostatic microglial markers such as P2ry12, Tmem119, Sall1, Gpr34, or Hexb. In addition to the DAM-like phenotype, d14 MCAo microglia showed enrichment of Gpnmb, Itgax, Csf1, Fn1, and Nceh1, supporting a chronic phagocytic, lipid-processing, and tissue-remodeling phenotype ^45^. Residual interferon-responsive genes (Ifit2, Ifit3, Ifi27l2a, and Bst2) further suggested that inflammatory signaling persists into the subacute-to-chronic phase ^32,44^.

Gene Ontology (GO) analysis of these microglial DEGs (SFig. 1d and STab. 4) pointed towards upregulated protein synthesis and energy production on d1, enhanced translation, activation of antimicrobial-like immune programs on d7 and d14 (“response to bacterium”) and retained enrichment of translational terms, together with broader immune effector terms such as “humoral immune response” and “cell killing” on d14. Finally, pseudotime inference suggested a transcriptional progression of microglia from sham-enriched, homeostatic-like states (clusters 2, 3, and 8) toward MCAo-associated activated states (SFig. 1c). Sham microglia from different time points largely overlapped in the embedding and occupied the earliest part of the inferred trajectory, consistent with a relatively conserved baseline microglial state across the sampled time points. Along the trajectory, cells were directed toward either d1 or d7/14 MCAo-enriched states, indicating a time-dependent shift in microglial transcriptional programs after ischemic injury. However, the trajectory of the day 1-to-day 7/14 transition was not observed in our data, indicating a non-linear remodeling of microglial states after stroke, in which acute inflammatory-responsive microglia and later phagocytic and lipid-remodeling microglia emerge as related but distinct transcriptional states.

### Transcriptionally distinct dendritic cell subsets are prominent interaction partners of microglia after ischemic stroke

Dendritic cells (DC) in homeostatic state reside mostly at the brain borders ^46^. They accumulate in the brain after ischemic injury, attracted to the parenchyma by microglia-secreted chemokines and cytokines, derived from blood and resident cell pool ^47^. Importantly, recent report highlighted the relevance of the intestine as a source of brain-infiltrating DCs in stroke ^48^. This further underscores the need for a detailed characterization of the gut-brain axis in ischemic brain injury.

The intercellular communication analysis (CellChat) of microglia interactions from the scRNA-seq dataset indicated that DCs are one of the main microglia interaction partners on all investigated timepoints (Fig. 2a-b; n= 25-40 CellChat outgoing interactions for specific DC subsets and timepoints). We identified robust signaling between microglia and cDC1, cDC2, moDC, and migDC on d1 and d14 after stroke, while d7 featured additionally interaction with pDCs (see SFig. 2 for ligand-receptor interactions microglia/dendritic cell subsets).

**Figure 2.**
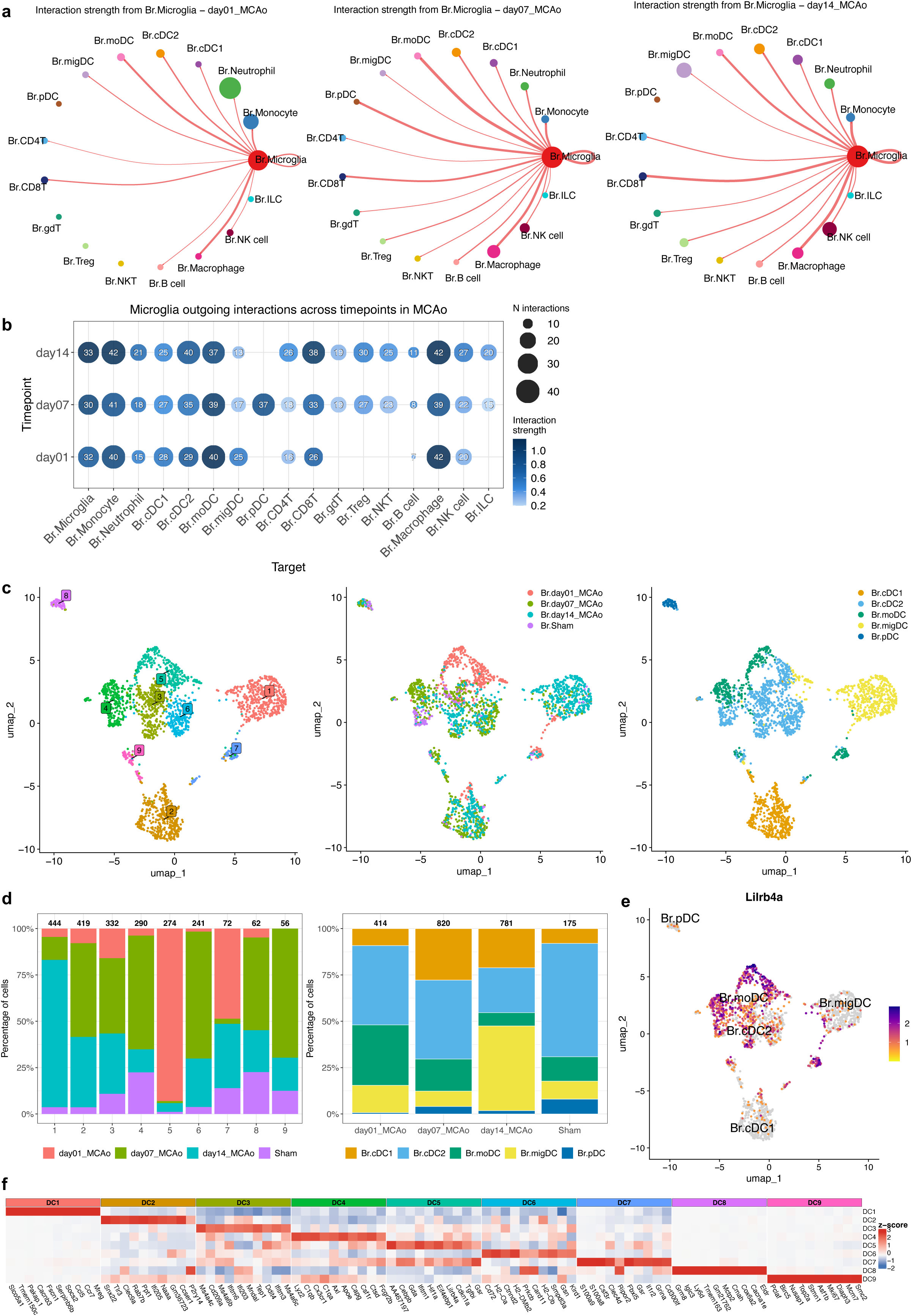
Dendritic cells (DCs) accumulation in the ischemic brain is accompanied by increased predicted interactions with microglia. **(a)** Outgoing communication networks of MCAo microglia in the ischemic brain across time points. CellChat circle plots showing inferred outgoing signaling from MCAo microglia at different post-stroke time points. Each node corresponds to a cell group and node sizes are proportional to the number of cells in each cell group. Red edges indicate outgoing communication from microglia to other cell populations and edge width represents the strength of the communication. The self-loop represents microglia autocrine communication. **(b)** Dot plot showing microglia outgoing interactions across timepoints in MCAo. Each dot represents the interaction from microglia to a target cell population at a given timepoint. Dot size indicates the number of interactions, and color intensity represents the aggregated interaction strength. Numbers inside the dots denote the corresponding interaction counts. (c) *left*: UMAP of DCs subset from brain reveals nine clusters; *middle* and *right*: UMAP of the same DCs colored by experimental groups (*left*) or colored by DC subpopulations (*right*). **(d)** Stacked bar plot showing the relative proportions of experimental groups across clusters (left) or of DC subpopulations (right) across experimental groups, numbers above the bars indicate the total number of cells in each group. **(e)** UMAP showing single-cell gene expression of Lilrb4a in DC subpopulations. Color scale represents log-normalized gene expression. **(f)** Heatmap of top 10 DEGs across brain DC clusters. For each DC cluster (DC1 – DC9), DEGs were identified by comparing that cluster with all other DCs. Top 10 upregulated genes in each cluster are shown in the heatmap. Heatmap columns represent genes and rows represent clusters. Color intensity indicates the scaled average expression level (z-score). Top 10 DEGs were selected based on |log2FC| ≥ log2(1.5), pct ≥ 0.75, and Bonferroni-adjusted p < 0.05, ranked by log2FC.

Closer examination of DCs transcriptional profiles revealed 9 distinct clusters in the brains of stroke and sham-operated mice, the vast majority of which was derived from stroke mice (Fig. 2c-d). On day 1 after MCAo, infiltrating DCs were dominated by conventional DCs type 2 (cDC2, identified based on canonical markers derived from literature, see Methods and STab. 1) and monocyte-derived DCs (moDCs). cDC2 and cDC1 were the main populations found in the brains on day 7 after stroke and day 14 featured migratory DCs as the dominant DC population. Besides that, we identified plasmacytoid DCs in the MCAo brains, with highest proportions on day 7 after stroke.

Considering the temporal resolution, on d1 after MCAo, DC cluster 5 was the most dominant one (cDC2 and moDC). Expression of Hif1a, Cebpb and Gsr in this cluster could reflect activation of cellular programs protecting from oxidative damage and reaction to infiltration of the ischemic environment ^49,50^. Although cDC2 are considered drivers of damaging inflammatory responses ^51^, a subset of these cells (50.2% of cDC2) expressed Lilrb4a known to mediate tolerogenic anti-inflammatory effects ^52,53^, while retaining the IFN-I responsiveness (expression of Ifitm1) (Fig. 2e-f; STab. 2). Day 7 after stroke featured DCs assigned to multiple clusters with most cells forming cluster 2 (cDC1) - expressing genes pointing towards their role in sensing damage signals (Tlr3, Clec9a, P2ry14) ^54^. Day 14 post- stroke had most cells situated in cluster 1 (migDCs), whose main role is to transport taken-up antigens to the lymph nodes. Indeed, presence of CNS antigens has been reported in draining lymph nodes after stroke ^55^. Besides their lineage markers, these cells expressed high levels of Ccl5, important for attracting other immune cells ^56^. Clusters 3 and 6 were populated mostly by cDC2 derived from d7 and d14 after stroke. Cluster 3 had high expression of IFN-response genes (Ifitm3, Ifitm6, Ifi203, Mndal) ^57^, cluster 6 genes related to antigen presentation and MHCII complex (H2-Oa, H2-Ob, H2-DMb2) ^58^ and inflammatory response (Card11 ^59^, Prkcb ^60^). Cluster 7 posed an interesting population with marked expression of genes related to the suppression of inflammation, including Cd300lf - recognizing phosphatidylserine on dying cells limiting DC inflammatory responses ^61^ or Il1r2, which is known to sequester IL-1 ^62^. Simultaneously, this cluster displayed upregulated expression of alarmins S100a8, S100a9 driving neuroinflammation ^63,64^ and Clec4d studied mostly in the context of responses to Mycobacterium ^65^ but also identified in myeloid cells in other stroke studies ^66^. Finally, cluster 4 was dominated by moDCs, cluster 8 by pDCs and cluster 9 presented a mix of DC subtypes with marked expression of cell proliferation markers (e.g. Mki67, Top2a, Birc5, Asf1b).

### Blood cells are minimally affected by transcriptomic changes after stroke, but engage in inflammatory programs in the brain

Parallel to the transcriptomic changes in brain-infiltrating leukocytes and microglia, we profiled transcriptomes from blood-isolated leukocytes of MCAo and sham-animals. Transcriptomic shifts when comparing MCAo to sham mice were limited in the 7 identified cell types (neutrophils, monocytes, DCs, NK cells, T cells, B cells and basophils) (Fig. 1c). Within these, the most prominent response was present in neutrophils on day 1 (151 DEGs, Bonferroni padj < 0.05 and |log2FC| >= 0.58). Top 10 (based on fold change, pct > 0.1) upregulated genes comprised: Chil3, Atp11a, Slc24a3, Pam, Per2, Gm36723, Flot2, Ahnak, Qsox1, and Ly6g (Fig. 3a-b; STab. 3), which points to a transcriptional shift toward a mobilized and migration-primed state. Notably, Chil3 (Ym1), induced after neutrophil activation in response to type 2 cytokines ^67^ has been previously detected in neutrophils infiltrating the brain after stroke ^68^. In parallel, Ahnak expression may indicate the presence of less mature or recently mobilized neutrophils, as AHNAK has been linked to immature neutrophil release from the bone marrow through regulation of the CXCR4 retention axis ^69^. Increased Flot2 further supports a migratory-ready phenotype, given the role of flotillins in neutrophil membrane-domain organization and uropod formation during cell polarization ^70^. Together, these changes suggest that acute stroke rapidly reshapes the circulating neutrophil compartment toward a mobilized, less mature and migration-primed state. By contrast, sham-enriched neutrophils showed higher expression of Cd101, a marker associated with mature circulating neutrophils in mice ^71^, supporting a relatively more homeostatic neutrophil phenotype in the sham condition.

**Figure 3.**
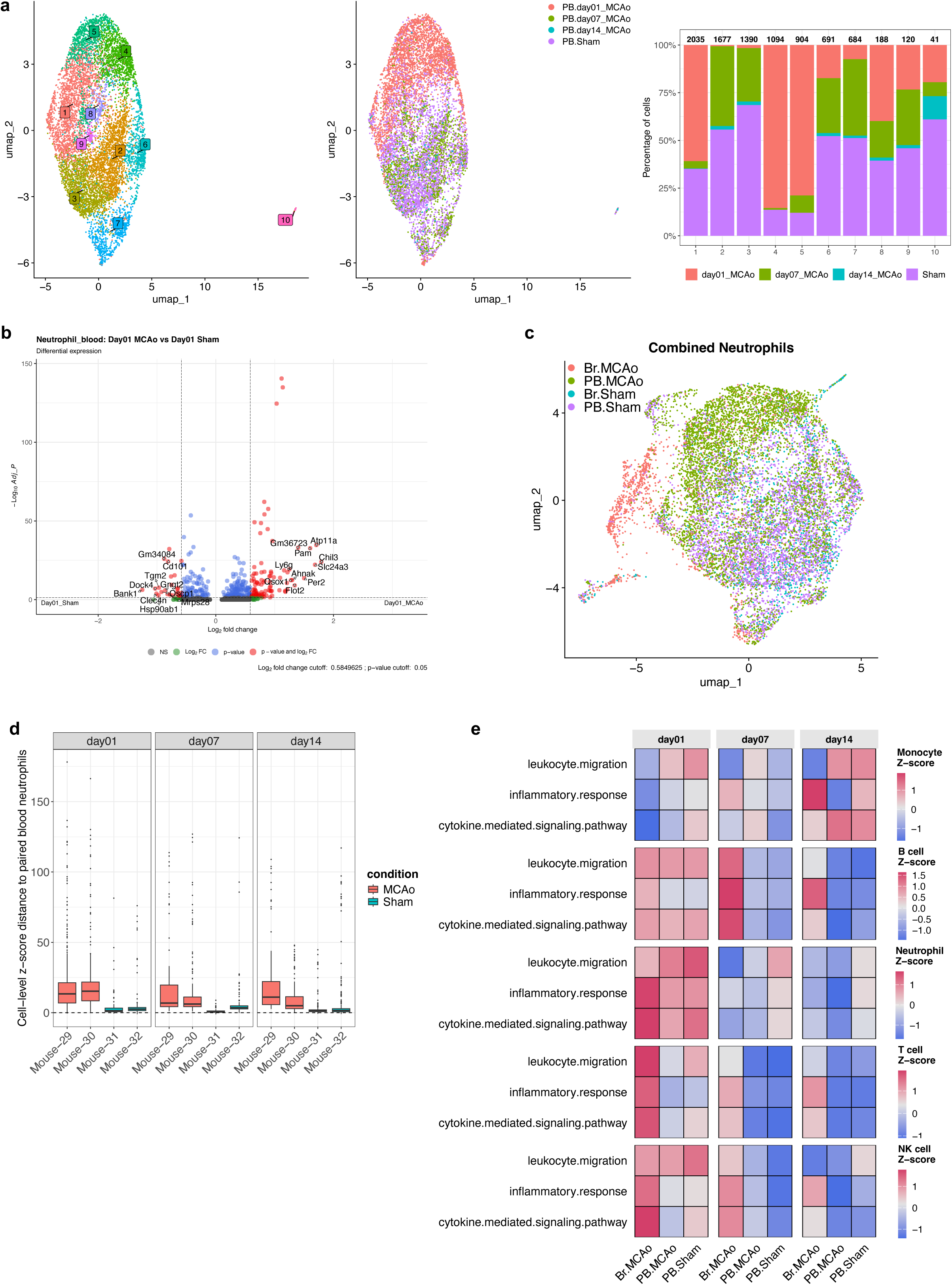
Blood leukocytes are minimally affected by transcriptomic changes after stroke while brain-infiltrating leukocytes engage in inflammatory programs. **(a)** *left*: UMAP of neutrophils subset from blood transcriptomes reveals ten clusters; *middle*: UMAP of the same neutrophils colored by experimental groups; *right*: Stacked bar plot showing the relative proportions of experimental groups across clusters, numbers above the bars indicate the total number of cells in each cluster. **(b)** Volcano plot of day 1 neutrophil DEGs in MCAo versus sham mice. Differential expression analysis was performed by comparing neutrophils from MCAo and sham groups. The top 10 DEGs (|log2FC| ≥ log2(1.5), pct ≥ 0.1, Bonferroni-adjusted p < 0.05, ranked by log2FC) are labelled in the condition in which they were enriched. **(c)** UMAP plots of neutrophils subset from the integrated brain and blood datasets. Brain- and blood-derived neutrophils are displayed in separate panels and colored according to experimental conditions. **(d)** Boxplots showing the cell-level z-score distance of brain-derived neutrophils to paired blood neutrophils from the same mouse. For each brain neutrophil, the mean distance to its five nearest blood-neutrophil neighbors was normalized to the within-blood distance distribution of the same mouse using the median and MAD. Lower z-scores indicate higher transcriptomic similarity to paired blood neutrophils and therefore a more blood-like profile. **(e)** Stacked heatmaps summarizing ssGSEA-based enrichment of inflammatory response, cytokine-mediated signaling, and leukocyte migration signatures in brain- and blood-derived leukocytes across experimental groups. For each immune cell type, normalized ssGSEA scores were averaged within each tissue-condition-timepoint group. Values are displayed as Z-scores calculated separately for each gene set across the displayed groups. Red indicates relatively higher enrichment, and blue indicates relatively lower enrichment.

Peripheral immune cells travel to the injured brain, which provides an inflammatory environment after stroke. We identified the following immune populations in the brain after ischemic injury: 1) neutrophils (7 clusters, with a relatively large population found in the sham brains SFig. 3), 2) monocytes, transcriptionally distinct on d1 after stroke from cells isolated in the subacute phase (7 clusters, SFig. 4) and 3) macrophages, prominent in the subacute phase after stroke (8 clusters, SFig. 5), 4) NK cells (5 clusters, SFig. 6), 5) T cells (CD8 dominating on d1, gamma delta T cells expanding on d7 after stroke and regulatory T cells, prominent on day 14 alongside CD4 and NKT cells, forming altogether 9 clusters - SFig. 7 for brain-infitrated and SFig. 8 for blood T cells) and 6) B cells (divided into 2 clusters, SFig. 9 and STab. 5 for DEGs across the timepoints). The unexpected abundance of neutrophils in sham brains (SFig. 3a), a population not typically resident in the healthy CNS, prompted us to examine whether these cells were truly brain-resident or reflected contamination from circulating blood. To address this, we integrated the brain and blood datasets and in the UMAP space and calculated the distances between brain-derived and blood derived neutrophils from stroke and sham mice (Fig. 3c). We hypothesized that the neutrophils in the sham group may be still contained within the brain vessels due to insufficient perfusion. Our analysis revealed greater distances from brain to blood neutrophils in the MCAo group (more dissimilar) as compared to the sham group, which would indicate similar transcriptomic states of sham neutrophils isolated from these two compartments, consistent with our hypothesis (Fig. 3d). This underscores an important challenge in MCAo studies and transcriptomic readouts relying on whole tissue preparations.

Furthermore, to get an insight into changes in the transcriptomic profiles of peripheral cells shaped by the neuroinflammatory milieu upon entry to the CNS, we compared blood and brain-infiltrated leukocytes using single-sample gene set enrichment analysis (ssGSEA). This method provides a score for all cells in a given population, assessing the activity of predefined pathways (GO terms). Given our hypothesis that the peripheral immune cells found in the brain have higher inflammatory activity than the cells found in blood, we used following GO terms in the ssGSEA analysis: inflammatory response (GO:0006954), cytokine mediated signaling pathway (GO:0019221), leukocyte migration (GO:0050900). We observed generally higher activity levels of these predefined gene sets in the brain for neutrophils, T cells, NK cells and B cells, especially on d1 after cerebral ischemia (Fig. 3e). Specifically “leukocyte migration” for monocytes was rather enriched in blood than in the brain, which aligns with the observations that monocytes migrate through blood to the ischemic brain where they differentiate into macrophages and dendritic cells ^72^.

### Stroke reshapes the intestinal proteome with early ileum immune mobilization and a day-7 peak of barrier and metabolic perturbation

The gut-brain axis is increasingly implicated in post-stroke pathology especially considering that the intestine is essentially an immune organ and home to the largest bacterial commensal community ^73^. It was recently shown that intestinal DCs migrate to the brain after ischemic stroke ^48^ and ca. 25% of brain infiltrating T cells are derived from the gut ^11^. To resolve the temporal dynamics and the magnitude of changes in the intestine after stroke, we performed proteomics from five intestinal segments (duodenum, jejunum, terminal ileum, caecum and proximal colon) and mesenteric lymph nodes (MLN) on timepoints 1, 7 and 14 after stroke or sham surgery (Fig. 4a). Within each tissue, timepoint dominated the variance over stroke condition (PERMANOVA R^2^=0.22-0.31 vs. 0.02-0.05), with day 1 samples (both MCAo and sham) diverging sharply from day 7/14 and highlighting the need for proper temporally matched controls in stroke experiments (SFig. 10). Within-timepoint stroke-vs-sham contrasts, however, mostly peaked later: 1 − π₀ (estimated proportion of truly differentially abundant proteins) reached 7–33% across compartments, highest at day 7 (Fig. 4a).

**Figure 4.**
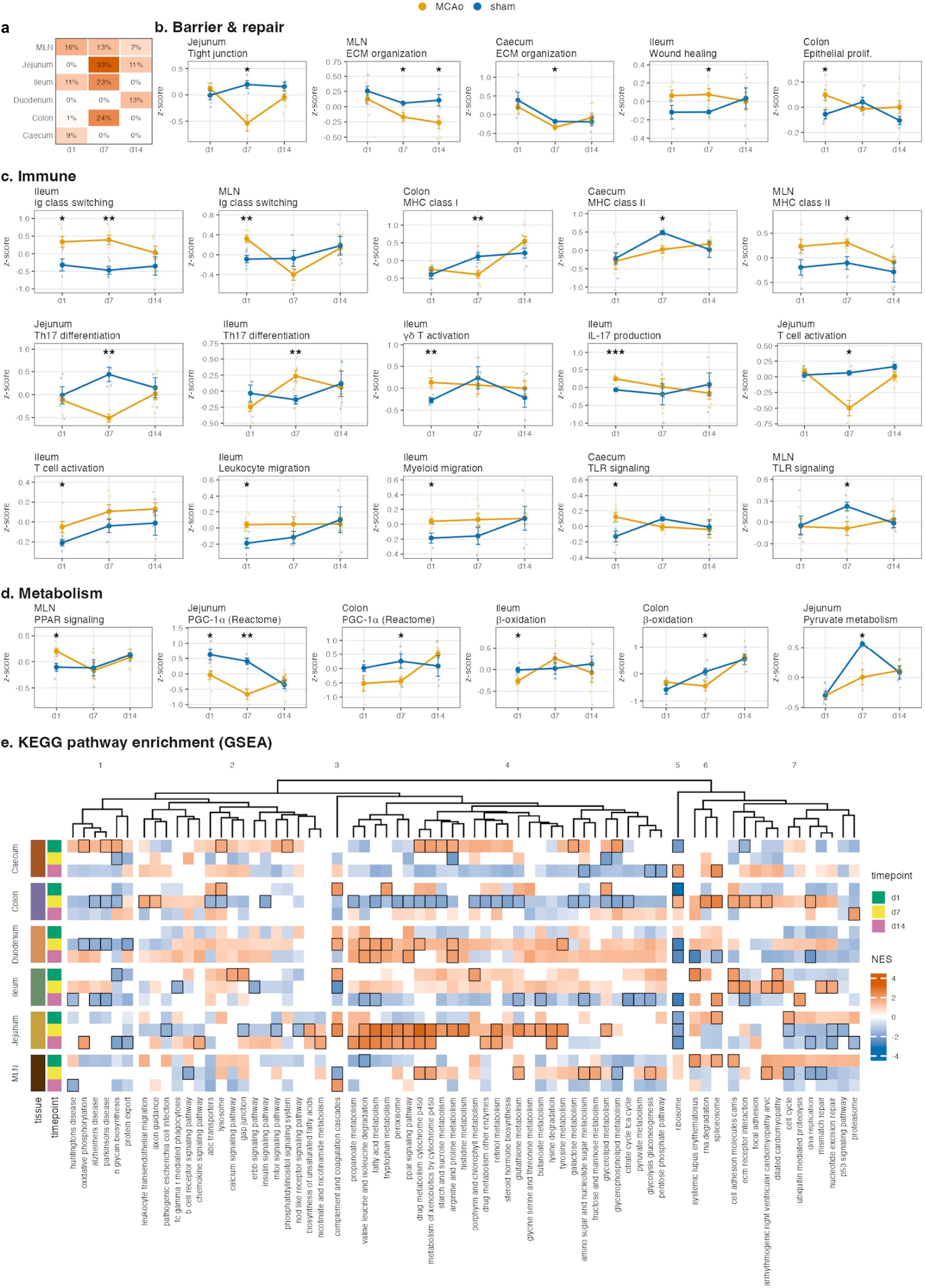
Intestinal proteomics reveals immune activation, barrier disruption and metabolic reprogramming after stroke, peaking at day 7. Bulk proteomic profiling of six intestinal tissues (caecum, colon, duodenum, ileum, jejunum, and mesenteric lymph nodes, MLN) from mice subjected to middle cerebral artery occlusion (MCAo, n=3–8 per tissue × timepoint) or sham surgery (n=3–5) at 1, 7, and 14 days post-stroke (176 samples total; 5,251-6,223 quantified protein-coding genes per tissue after gene-symbol mapping and QC). Surrogate variables capturing sacrifice-date and batch effects were estimated by SVA (2–5 surrogate variables per tissue) and removed before downstream analysis. **(a)** Proportion of proteins with non-null effects (1 − π₀, estimated proportion of truly differentially abundant proteins) estimated from the p-value distribution of MCAo vs sham differential expression (limma, full model adjusting for SVs) per tissue and timepoint. π₀ is a descriptive summary of the global p-value distribution and not itself a test statistic; signal peaks at day 7, notably in jejunum (33%), colon (24%), and ileum (23%). **(b-d)** Hypothesis-driven gene-set analysis of pre-defined biological processes spanning barrier and repair **(b)**, immune **(c)**, and metabolic **(d)** programs. Gene sets were taken mainly from GO:BP via the msigdbr R package, org.Mm.eg.db (GO:0030198) and PGC-1α activation from Reactome. For each gene set, expression values were z-scored per gene across all samples within a tissue, averaged into a per-sample score, and compared between MCAo and sham within each timepoint by two-sided Welch’s t-test. Each panel shows the mean ± SEM trajectory of MCAo (orange) and sham (blue) across timepoints in tissues where a significant difference was detected (nominal p < 0.05 on SVA-corrected data); individual sample scores are shown as jittered points. Asterisks indicate per-timepoint significance: *p < 0.05, **p < 0.01, ***p < 0.001. Nominal p-values are reported because the gene sets are pre-specified hypotheses, not a discovery scan; exact p-values for every tissue × timepoint × gene-set combination are provided in STab. 6. **(e)** Gene set enrichment analysis (GSEA) of 186 KEGG pathways across all tissue-by-timepoint combinations, ranked by limma moderated t-statistic on SVA-corrected data. After collapsing redundant gene sets, non-redundant pathways significant in at least one contrast (BH-adjusted p < 0.05) are shown. An entry-level direction-concordance filter was applied: each entry (tissue x timepoint x pathway) shows the SVA-corrected NES only where the sign of enrichment matches that of the parallel uncorrected analysis; entries whose direction is reversed by SVA correction are masked (white). 69 of the 72 pathways had at least one direction-concordant cell and are shown; 364 of 1,242 cells (29%) are masked. Pathways are hierarchically clustered (Euclidean distance, complete linkage). Rows are tissue × timepoint combinations annotated by timepoint color (d1, d7, d14); columns are pathways split into seven clusters. Bordered cells mark significant enrichment (BH-adjusted p < 0.05 in the SVA corrected data and direction-concordant with uncorrected data).

We next used a hypothesis-driven approach to test pre-defined biological processes spanning barrier and repair, immune, and metabolic programs across all sections and timepoints, using curated GO: BP and Reactome gene sets (STab. 6). We first asked whether stroke compromises the gut barrier ^11,14,17^. Gene-set z-scores for tight junction organization (jejunum, d7) and ECM organization (caecum, d7) were coordinately decreased in MCAo vs sham (Fig. 4b), while ECM remodeling also extended to the MLN (d7, d14), reflecting sustained tissue remodeling in the gut-draining lymphoid compartment. Two repair-associated programs accompanied this barrier disruption: wound healing in ileum at d7 and epithelial cell proliferation in colon at d1, consistent with active mucosal repair (Fig. 4b).

We next tested immune hypotheses motivated by reports of gut-to-brain T-cell migration ^11,18^, suppression of local immune responses ^74^, and the immune activation observed in our scRNA-seq data. The ileum emerged as the principal site of early gut-immune mobilization at day 1, with a coordinated set of stroke-specific increases: IL-17 production, γδ T cell activation, T cell activation, leukocyte migration, and concordant myeloid migration (Fig. 4c). By day 7, the ileum signature shifted toward Th17 differentiation and immunoglobulin class switching, indicating a progression from γδ T-cell early IL-17 production at d1 toward Th17-lineage and humoral responses at d7. In contrast, the jejunum showed opposing decreases in Th17 differentiation and T cell activation at day 7 (Fig. 4c), highlighting compartment-specific divergence within the small intestine. The MLN showed early Ig class switching (d1), MHC class II presentation increase (d7), consistent with its role as an integrative node between gut-derived signals and adaptive priming. Antigen presentation diverged across compartments: MHC class I decreased in colon (d7), MHC class II decreased in caecum (d7) but increased in MLN (d7), and TLR signaling increased in caecum (d1) but decreased in MLN (d7), indicating tissue-specific reorganization rather than a uniform program. Given that the small intestine hosts the largest mucosal T-cell compartment and is a source of gut-derived peripheral T cells, the temporal sequence of γδ T/IL-17 (d1) to Th17/Ig class switch (d7) to sustained MLN remodeling (d14) positions the ileum as a candidate peripheral source for the brain γδ T-cell expansion observed at day 7 and the evolving T-cell composition at day 14 (SFig. 7).

We next tested metabolic hypotheses linked to host energy states and availability of microbial short chain fatty acids (SCFAs). In a healthy gut, colonocytes rely on SCFAs as their primary energy source used in β-oxidation; the resulting oxygen consumption keeps the lumen hypoxic and sustains the anaerobic microbiome ^75^. This equilibrium is disturbed in intestinal inflammation and/or intestinal ischemia, which have been previously reported after stroke ^12,76^. Activation of PGC-1α, a master regulator of mitochondrial biogenesis and fatty-acid oxidation ^77^, was decreased in jejunum (d1, d7) and colon (d7), with parallel decreases in β-oxidation (ileum d1, colon d7) and pyruvate metabolism (jejunum d7) and an isolated PPAR signaling increase in MLN at d1 (Fig. 4d). Together, these point to a switch in enterocyte metabolic programs at d7 toward reduced oxidative capacity, predicted to raise lumen oxygenation and to enable facultative-anaerobe expansion ^78^.

To complement the hypothesis-driven tests with an unbiased pathway-level view, we performed a gene set enrichment analysis (GSEA) on KEGG pathways across all tissue x timepoint MCAo-vs-sham contrasts, ranking proteins by limma moderated t-statistic on SVA- corrected data, leveraging the complete distribution of quantitative data without imposing an arbitrary significance cutoff (Fig. 4e). To prevent batch-correction artefacts, an entry-level direction-concordance filter was applied (see Fig. 4 caption): BH-significant pathways (adjusted p < 0.05 in at least one contrast in SVA corrected data) that are direction-concordant are retained. We noted an opposite direction of pathway responses in d7 jejunum and colon (Fig. 4e). Jejunum d7 was positively enriched for many metabolic and detoxification pathways: amino-acid degradation (BCAA, tryptophan, histidine, tyrosine, arginine/proline, lysine, β-alanine, glycine/serine/threonine), lipid and fatty-acid handling (fatty-acid metabolism, peroxisome, PPAR signaling, butanoate, glycerolipid, unsaturated FA biosynthesis), xenobiotic and drug metabolism (cytochrome P450 drug metabolism, xenobiotics by cytochrome P450; retinol metabolism), glutathione metabolism, starch and sucrose metabolism, and complement and coagulation cascades.

Colon d7 mirrored almost the same set in the opposite direction (fatty-acid handling, xenobiotic metabolism, glutathione, starch and sucrose), with additional negatively enriched pathways: steroid hormone biosynthesis, propanoate, galactose, amino-sugar and nucleotide sugar metabolism, insulin signaling, TCA cycle, glycerophospholipid, N-glycan biosynthesis, and porphyrin and chlorophyll metabolism. Counter to this, colon d7 also showed positive enrichment of biosynthetic and tissue-remodeling pathways - spliceosome, RNA degradation, DNA replication, ECM-receptor interaction, focal adhesion, cell adhesion molecules, leukocyte transendothelial migration, and ribosome - indicating a shift toward proliferative and immune-traffic programs.

On day 1, caecum and ileum carried the strongest pathway-level responses. Caecum d1 showed positive enrichment of carbohydrate and energy metabolism (glycerophospholipid, galactose, oxidative phosphorylation, lysosome) and xenobiotic and drug metabolism, alongside negative enrichment of ribosome, and ECM-receptor interaction. Ileum d1 was positively enriched for complement and coagulation cascades, gap junction, calcium signaling, porphyrin and chlorophyll metabolism, cell adhesion molecules, and immune-related pathways (systemic lupus erythematosus), consistent with the early immune-mobilization phenotype seen in panel c (γδ/IL-17). Two pathways stood out across the dataset for breadth and magnitude: ribosome was strongly downregulated at d1 with compartment-specific recovery by d7 or d14, suggesting an early translational-machinery suppression followed by tissue-specific rebound; and complement and coagulation cascades were broadly upregulated across compartments and timepoints, consistent with sustained innate-immune activation through the post-stroke window.

Taken together, the proteomic response segregates into three temporally distinct modules: (i) an early ileum-centered γδ T/IL-17 immune mobilization at day 1, distinguishable from the surgical-stress background that dominates whole-tissue variance at this timepoint; (ii) a day-7 peak of stroke-specific changes across compartments, coupling barrier disruption markers (jejunum TJ, caecum/MLN ECM) with compartment-specific metabolic reorganization, coordinated decreases in PGC-1α–driven enterocyte oxidative programs in jejunum and colon (Fig. 4c-e), and Th17/humoral maturation in ileum and MLN; (iii) sustained matrix remodeling in the gut-draining lymph nodes through day 14. This temporal architecture motivates examining the gut-microbial response over the same window, where we anticipate stroke-specific microbial changes to emerge alongside the host-metabolic alterations at day 7.

### Stroke-specific dysbiosis coupled with gene expression changes emerges beyond the surgical stress window and features an expansion of facultative anaerobes

We aimed to characterize stroke-induced changes in the gut microbiome using metatranscriptomics (longitudinal collection of samples: one day before the surgery (d-1), d1, d3, d7 and d14 thereafter) and metagenomics (d14). To the best of our knowledge, changes in the gut microbiota transcriptome after ischemic stroke have not been reported until now and our dataset provides unique insights into longitudinal metatranscriptomic profiles. We found species abundance varying over time (Fig. 5a) and some of them showed different abundances associated with condition and time. *Dubosiella muris* (MAG024) showed negative association with timepoint d3, d7 and stroke; Lachnospiraceae (MAG041) showed positive association with d1 and d3, but negative association with stroke. Finally, *Ligilactobacillus murinus* (MAG050) was positively associated with d1, d3, d7 and d14 and also with stroke (Fig. 5b). The dynamics of the microbiome response showed a complex and multilayered pattern consisting of heterogeneity between replicates, effect of housing/cage assignment (groups were housed separately to avoid the confounding microbiome transfer effect ^13^), surgery and finally stroke. Interestingly, time had the strongest effect across all analyses, highlighting the importance of longitudinal microbiome sampling rather than single-endpoint measurements. In order to disentangle the different factors contributing to microbiome dynamics, we compared distances between samples over time using Aitchison distance, which explicitly accounts for the compositionality of the abundance data. We observed that samples from stroke and sham-operated mice (between sample distances) became more similar after surgery while dissimilarity increased afterwards to approximately the same levels as the baseline (Fig. 5c). Similarly, the variability of samples from the same condition (within sample distances) decreased with surgery and increased again later (SFig. 11a). Both comparisons showed a significant effect of time when correcting for cages and replicates. Interestingly, we could detect a clear difference between MCAo and sham animals when we followed individual trajectories and compared the distances of each microbial community to its baseline (Fig. 5d). Here, microbiomes of stroke mice diverged with higher distances compared to day -1, specifically starting from day 3. The effect was still visible on day 14, indicating a lasting perturbation that is not seen in sham mice (linear mixed effect model with cage and replicate as random effects, p-value of stroke condition effect 0.026 calculated using ANOVA with Satterthwaite’s method). In summary, comparing the similarity of samples revealed variability between replicates that initially decreased after surgery and increased later again, whereby particularly stroke microbiomes diverged significantly from before treatment conditions.

**Figure 5:**
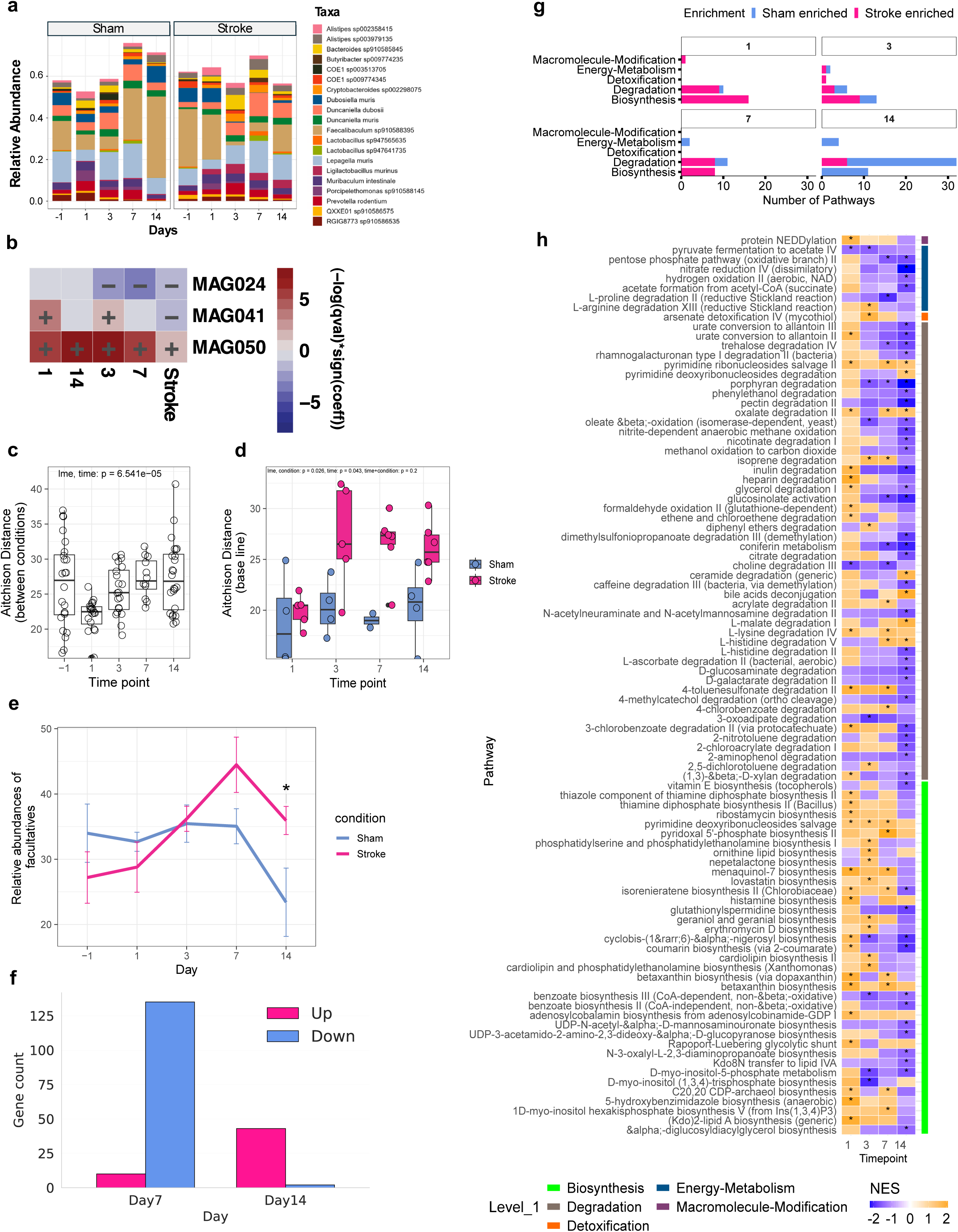
**Shifts in gut microbial diversity and pathways from metatranscriptomics following stroke**. **(a)** Mean relative abundance of taxa with at least 0.1 relative abundance at each timepoint. **(b)** Differentially abundant species detected by MaAsLin2 showing positive and negative associations with condition and time. Only species with significant association with condition (stroke or sham) were selected, i.e.: *Dubosiella muris* (MAG024), *Lachnospiraceae* (MAG041; taxonomy resolved only on family level), *Ligilactobacillus murinus* (MAG050). **c)** Dissimilarity between samples from different conditions measured by Aitchison distance. **(d)** Dissimilarity of samples compared to baseline (day -1) for each animal. **(e)** Change in the relative abundance of facultative anaerobes in stroke and sham across timepoints. **(f)** Gene Count of differentially expressed genes between conditions for each timepoint obtained by dream (differential expression for repeated measures). **(g)** Count of sham enriched and stroke enriched significant pathways between conditions for each timepoint summarized by subsystems (Level-1 Metacyc hierarchy) obtained by GSEA **(h)** sham enriched and stroke enriched significant pathways between conditions for each timepoint obtained by GSEA. Heatmap displays normalized enrichment scores (NES) from gene set enrichment analysis (GSEA) of non-redundant MetaCyc pathways at days 1, 3, 7, and 14 following stroke. Color intensity reflects NES magnitude with warm colors (orange) indicate stroke-enriched pathways (NES > 0 & padj < 0.05) and cool colors (purple) indicate sham-enriched pathways (NES < 0 & padj < 0.05). Asterisks denote pathways passing the significance threshold (padj < 0.05). Pathway redundancy was minimized using the collapse function from the fgsea package. Right-side color bar indicates MetaCyc level-1 functional hierarchy classification.

We next asked if diverging microbial communities in stroke could be linked to changes in microbial functional groups. Microbiome dysbiosis in the gut could often be attributed to the overgrowth of facultative anaerobes exploiting inflammatory conditions ^79^. Therefore, we predicted the oxygen preferences based on the genomes assembled from metagenomic sequencing and found an increase of facultative anaerobic bacteria in MCAo animals, starting after d3 (Fig. 5e, linear mixed effect model with cage and replicate as random effects, p-value of stroke condition effect 0.036 calculated using ANOVA with Satterthwaite’s method). Furthermore, to characterize the transcriptional response in the microbiome, we performed differential expression analysis of metatranscriptomic data using Dream, a differential expression framework for repeated measurements ^80^.

Leveraging the advantage of longitudinal collection and d-1 sample, we performed 3 types of analysis: 1) case I = within-condition temporal contrasts, comparing samples against their respective baselines; 2) case II = cross sectional contrasts, comparing stroke and sham samples at the same timepoints; 3) case III = interaction contrasts assessing changes relative to baseline and between conditions (STab. 7, SFig. 11a). In case I, the stroke group showed substantial transcriptional changes at all timepoints relative to day -1 with larger effects detected on d3 and d7. In contrast, within sham samples, we observed notable changes at d1 and d3 with minimal to no changes at d7 and d14. This stark difference between the two conditions suggests a stronger and more sustained stroke-associated response at later timepoints. Cross-sectional comparison between stroke and sham mice at individual timepoints (case II) revealed most DEGs on d7 (mostly downregulated in stroke animals) followed by d14 (majority of DEGs being upregulated, Fig. 5f). This observation - most pronounced differences between stroke and sham mice emerging on day 7 is consistent with host gut proteomic profiling. No significant differences were detected on d1, or d3, indicating that the two groups were transcriptionally comparable with possible dominance of surgery effect at early timepoints and emergence of transcriptional changes later. Interaction contrasts (case III), did not yield significantly DEGs except d7 with only 5 genes after FDR correction. The convergent evidence from within-stroke temporal contrasts and cross-sectional comparisons at days 7 and 14 collectively supports a stroke-specific transcriptional response that seems to intensify at later timepoints.

To understand how stroke reshapes the functional capacity of the gut microbiome, we performed GSEA with MetaCyc pathway annotations at Levels 1 and Level 2 hierarchies, tracking changes across four timepoints (days 1, 3, 7, and 14) post-stroke. The number of significant pathways at each timepoint and comparison is summarized in the STab. 7. Leveraging the MetaCyc hierarchy allowed us to examine pathways relative activity at both broad and finer levels of resolution using normalized enrichment scores (NES) highlighting temporal patterns spanning five major categories: biosynthesis, degradation, energy metabolism, macromolecule modification, and detoxification, and 46 sub-major categories (Level-2 MetaCyc hierarchy, STab. 7). When stroke and sham microbiomes were directly compared at each timepoint (Case II), a consistent pattern emerged: stroke-associated communities were skewed toward biosynthesis at early timepoints, while sham communities showed stronger enrichment of degradation and energy metabolism pathways over time (Fig. 5g). We observed the same trend across the other two comparisons as well. Within the biosynthesis category, sham-enriched pathways showed a compositional shift over time. Early timepoints were characterized by enrichment of secondary metabolite, cofactor, and lipid biosynthesis pathways in stroke, which were progressively replaced at later timepoints by the enrichment of aromatic compound and carbohydrate biosynthesis pathways in sham. In the degradation category, sham-enriched degradation pathways contributed majorly by aromatic compounds degradation, carbohydrates degradation, carboxylates degradation, non-carbon nutrients, secondary metabolite degradation, showed a consistent that increased consistently across all timepoints, while stroke-associated degradation signals e.g. from including amino acid degradation, fatty acid/lipid-degradation, carboxylates degradation, carbohydrates degradation were variable and directionally mixed. Energy metabolism followed a similar pattern, with sham-associated enrichment accumulating steadily from day 3 through day 14 with pathways mostly primarily driven by chemoautotrophic energy metabolism, pentose phosphate cycle and fermentation. In contrast, stroke-associated enrichment of the energy metabolism pathway stemmed primarily from fermentation and was largely confined day 3 (Fig. 5h, SFig. 11b).

## Discussion

Our study provides comprehensive immune transcriptomic profiling post-stroke - day 1, 7 and 14 after cerebral ischemia coupled with proteomics of five intestinal segments and mesenteric lymph nodes alongside longitudinal metatranscriptomic analysis (one day before MCAo/sham, day 1, 3, 7 and 14 thereafter) and metagenomics (day 14) derived from the gut microbiome. We show that whereas the immune system responds rapidly to the injury in the brain as evident by transcriptomic changes in microglia and brain-infiltrating immune cells, the stroke-specific changes in the periphery emerge clearly at later timepoints. Profiling multiple conditions within the same organism constitutes a major strength of this study, as it circumvents inter-individual variability and enables a more precise characterization of stroke-specific signatures. Furthermore, this study provides the first metatranscriptomic dataset from the experimental stroke model and highlights the need for time-matched controls in stroke studies as well as proper controlling for batch and cage effects, well known in microbiome research.

Analyzing the post-stroke brain immune landscape, we focused on microglia and dendritic cells as hubs of post-stroke immune events. We identified 9 major microglia clusters based on their transcriptomic profiles. On day 1 after cerebral ischemia, the microglia showed upregulation of genes associated with active inflammatory signaling and phagocytosis. Day 7 and 14 featured a switch to the typical DAM programs, reparative lipid-handling microglia and upregulation of the IFN-I - activated genes. Among these, Ifi27l2a with increased expression on both subacute timepoints after MCAo in our dataset, has been described as the master regulator of IFN programs after stroke and in aging ^32^. Interestingly, we observed cluster 1 as a prominent microglia subtype co-expressing DAM markers, MHC-I molecules and IFN-I stimulated genes emerging on day 7. This cluster expressed also many markers of the Ch25h+ subset identified by Zhang et al. ^43^ but together with Oasl genes, which diverges from the suggested division into protective Ch25+ and detrimental Oasl+ microglia after stroke ^43^ and warrants further investigations. GO enrichment analysis revealed up-regulation of microglial gene pathways involved in antigen presentation alongside “response to bacterium”. Intriguingly, recent reports have raised the possibility of bacterial translocation from the gut to the brain after experimental stroke ^81,82^. The genes underlying this GO term are relevant for two scenarios - sterile inflammation and immune activation in response to bacteria. While the majority of available literature supports the former explanation, we did not test for presence of bacteria or bacterial components in other organs than the gut in our study.

Our snRNA-seq analyses indicate an intense microglia-dendritic cell signaling on all investigated time-points after stroke. Although microglia can act as antigen-presenting cells, it is known that DCs surpass microglia in this function ^47^. Microglia, as brain-resident macrophages, are the first responders to the ischemic injury coordinating further immune cascade. They are responsible for early sensing and clearance of the damaged area, while specialized immune jobs like antigen presentation are handed over to DCs when they arrive at the ischemic injury site. We confirmed that cDC2 is the dominant subset on day 1 and the presence of cDC1, pDCs, moDCs and migDCs in the post-stroke brain as described by others ^6,51^. Interestingly, most moDC and a large subset of cDC2 found in the brains on day 1 after stroke, expressed Lilrb4a, which is a known inducer of tolerogenic states ^83^. In general, cDC2 are regarded as a detrimental DC subset attracting γδ T cells to the injury site driving post-stroke neuroinflammation ^47^. Therefore, cDC2 upregulation of Lilrb4a expression in injury states could provide a safety mechanism protecting from induction of excessive inflammatory response ^52^. Moreover, pDCs seem particularly interesting in the microglia-DC interactions, as these DCs are specialized IFN producers ^84^. IFN-I can be released by many cells after stroke, with some reports pointing to the central role of Th1 cells in IFN-I mediated signaling ^85^, however pDCs have not been studied in this context so far. Although IFN-I responsive microglia are generally considered detrimental in CNS injury ^30,86^, one study indicates that pDC depletion is worsening stroke outcome in the experimental model ^87^. Other reports show that exogenous IFN- α and IFN-β delivery is beneficial after stroke ^88^, underlining the need for an in-depth characterization of IFN-I responses post stroke ^35^.

We have not identified pronounced transcriptomic shifts in the blood peripheral cells, although it is known that ischemic stroke induces an immunosuppressive state during the first days after brain injury ^8,9^. Our analysis timepoints are, however, at the borders of the time when immunosuppression is most pronounced ^8,89^ and the interpretation of scRNA-seq is further complicated by limited availability of blood transcriptomic datasets from experimental stroke models ^90^. When comparing blood and brain-infiltrating cells, our data indicate that peripheral blood cells activate inflammatory programs upon entry to the brain which is supported by the concept of transcriptomic switch / “transcriptomic divergence” ^6^ in these cells.

A central and initially counter-intuitive observation is that stroke-specific peripheral signatures emerge in the subacute window (days 7 to 14) rather than on day 1, in apparent tension with prior reports of rapid post-stroke dysbiosis ^11,12^ and drastic effects on the gut, including large shifts in gene expression in the colon ^13,91^. In our experiments, time-matched sham controls together with correction for batch and cage effects absorb a substantial early surgical-stress signal that is shared between MCAo and sham, leaving within-timepoint stroke-vs-sham contrasts largely null at day 1. However, similarly to previous findings ^12,92^, day 7 in our study features an expansion of facultative anaerobes, although we have not investigated its functional consequences (e.g. previously suggested detrimental impact on the outcome ^12^). This phenomenon links the microbiome readouts to proteomics. In intestinal homeostasis, colonocytes use SCFAs, particularly butyrate, as energy source, consuming high oxygen levels in the process of β-oxidation, maintaining an anoxic gut environment ^75^. Without butyrate, colonocytes switch to anaerobic glycolysis, which leads to a leakage of oxygen to the gut ^75^ and enables an increase in facultative anaerobes.

Furthermore, our proteomics analysis points towards day 7 after stroke as a critical timepoint for changes in the tight junction and ECM markers (which could be connected to alterations in permeability). Gut permeability changes after stroke have been reported previously, although occurring again rather on early timepoints ^11,14,17^. Our proteomic dataset highlights the gut as an active immune organ, with marked compartmentalization as specific changes are section-and time-point specific, which warrants further investigations. When it comes to the divergence between the proteomic changes in jejunum vs. colon, we interpret these differences as linked to compartment-specific biology. The small intestine, including jejunum is the major site of nutrient absorption, features less microbes and more aerobic environment. In the colon, the oxygen abundance is significantly lower with increasing microbial mass ^93^. We propose therefore that the divergence between jejunal and colon responses may be connected to the changes in the microbiota, which we observe in the colon compartment. To fully test this hypothesis, sampling from multiple gut compartments is needed in future studies. Another possible explanation may be linked to immune function of the jejunum carrying an active T-cell signature (not present in the colon). As activated T cells run on glutaminolysis, leucine-driven mTOR, and amino-acid catabolism ^94^, the jejunal enrichment could reflect changes in immune cells rather than enterocytes.

Changes in the gut proteome paralleled alterations in microbial gene expression, where we found most DEGs at later time points. Positive enrichment of multiple pathways already on day 1 in the GSEA may be explained by higher sensitivity of this analysis to identify subtle changes ^95^. Specifically, we found biosynthesis pathways enriched in stroke animals on early timepoints (most prominent on day 1). This could be connected to changes in the food intake in the first days after MCAo and the compensatory microbiome response. Reduced food intake (especially on early timepoints) and weight loss in experimental stroke animals are well known phenomena. The weight loss, however, is not only a result of decrease in food consumption, but involves changes in overall metabolism and mobilization of fat and muscle reservoirs ^96,97^. It has been also suggested that the weight loss is necessary for stroke recovery in the experimental model and linked to corticosterone release crucial in coordinating post-stroke immune responses ^98^.

Considering specific MetaCyc pathways, two connected to the availability of SCFAs showed negative enrichment in stroke animals: “pyruvate fermentation to acetate IV” and “acetate formation from acetyl-CoA” (for this pathway, except day 1). SCFAs are not only energy sources but also mediators of immune effects, including modulation of DCs and T cells functions ^99^. In general, supplementation of SCFAs after stroke leads to beneficial outcomes through fine-tuning of lymphocytes and microglia responses ultimately leading to reduced neuroinflammation ^76,100^. Protective SCFA effects on the gut and intestinal barrier have been also proposed as relevant in stroke ^82^. Further, we noted positive enrichment of bile acids deconjugation in stroke mice (d14). This process is an important step in production of secondary bile acids, which mediate protective effects after stroke ^101^. Finally, the histamine biosynthesis pathway was positively enriched in stroke animals (significantly on d1). Histamine is a neuroactive and immunomodulatory metabolite, with different actions depending on the target receptors. It can activate proinflammatory signaling (H1 and H4 receptors), e.g. through mast cell degranulation or cytokine secretion from invariant NKT cells ^102^ but also suppress inflammation and promote Treg polarization (H2 receptor ^102^, which may be relevant for shaping the outcome after stroke as Tregs are considered generally protective in experimental stroke models ^103^). The neuromodulatory actions on enteric neurons would be mediated over the H3 receptor ^102^.

Finally, we did not observe massive changes in the microbial community composition after stroke, although stroke animals diverged more from the baseline composition than sham animals. *Dubosiella muris* and Lachnospiraceae were negatively associated with stroke and *Ligilactobacillus murinus* was positively associated with stroke. *L.murinus* is known for its anti-inflammatory properties ^104^, supporting the intestinal barrier ^105^ and mediating suppression of macrophage cell death ^106^. We hypothesize that an increase in the abundance of this bacterium may be a compensatory response after stroke.

Altogether, this report provides temporally resolved multi-omic characterization of host immune response to ischemic stroke, highlights day 7 as a time-point of altered gut proteome and gut microbiome function and underlines the need for inclusion of matched sham controls.

Limitations:

Despite the insights provided, our study is not without limitations. Only young, male mice were included in our investigations to minimize heterogeneity between experimental animals. Further tests should include mice of both sexes and ideally of several age categories. As this study focuses on an exploratory analysis, we performed scRNA sequencing with n=2 per group and timepoint. Blood collection on day 14 suffered from technical problems and only a limited number of cells has been recovered from this timepoint (SFig. 1b). Finally, future omic analyses should also include Peyer’s Patches as recent reports show that this immune gut compartment is particularly affected after ischemic stroke ^74,91^.

## Materials and methods

### Animals

Adult (11-14 weeks old, at the start of the experiment) male C57Bl/6J mice (Janvier Laboratories, Le Genest-Saint-Isle, France) were used in this study. All mice were housed at the FLI animal facility in individually ventilated cages with a 12 h light/dark cycle and ad libitum access to food and water. All animal experiments were performed in accordance with the ARRIVE guidelines, European Community Council directives 86/609/EEC and German national laws, and were approved by the local authority (Thüringer Landesamt für Verbraucherschutz). One cohort of n=40 animals was included in this study with subsampling for specific scientific outcomes: n=2 mice per condition per timepoint (total 12 mice) were used in the scRNA sequencing experiment. Material for intestinal proteome analyses was sampled from all mice and n=3-8 per compartment, condition and timepoint were included in the analyses. Stool samples from n=2-6 mice per condition and timepoint were included in the metagenomics/metatranscriptomics readouts. Reaching humane endpoints, failure of collecting the samples (e.g. stool samples) and significantly lower protein yield when compared to all other samples in proteomics were used as exclusion criteria in this study.

### Induction of experimental stroke and organ sampling

Mice were subjected to 45 min middle cerebral artery occlusion (MCAo) as described in ^107^ followed by reperfusion. Full sham surgery (including insertion of the filament, followed by immediate withdrawal) was implemented in the control group. Perioperative analgesia was provided by administration of metamizol in drinking water ad libitum (200 mg/kg one day before surgery and switching to 100 mg/kg on the day of surgery continued until day 3 inclusive). All mice were provided wet mash food and pellets at the cage floor. Subcohorts of animals were sacrificed at 1, 7 or 14 days after surgery. Considering the potential of microbiome transfer within the cage, MCAo animals were housed separately from sham-operated mice. Fecal samples were collected 1 day before surgery (day -1), and 1, 3, 7, and 14 days after surgery at the same time of the day. Fresh fecal pellets were collected directly from mice and stored at −80 °C until further analysis. At the endpoint, mice were anesthetized with a mixture of ketamine and xylazine (ip. injection) and blood was collected from *vena cava caudalis* into EDTA tubes and kept on ice until leukocyte isolation. Afterwards, cardiac perfusion was performed, and brain and intestines sampled for further analyses. Ipsilateral hemispheres were separated from the cerebellum and olfactory bulb, rinsed by Dulbecco’s Balanced Salt Solution without calcium and magnesium (DPBS, Thermo Scientific) and kept on ice for a short time until cell isolation. The whole intestine was collected cutting behind the pylorus and at the rectum and subsequently sectioned into five regions of interest (1 cm of tissue sampled): proximal duodenum, proximal jejunum, distal ileum, tip of the caecum, proximal colon. The tissues were flushed with PBS and frozen in liquid nitrogen for further processing. Mesenteric lymph nodes were collected separately for further analysis.

### Preparation of single cell suspensions

For isolation of blood leukocytes, blood from each mice was processed according to the protocol of leukocytes isolation for scRNA-sequencing (CG000392, Rev A, 10X Genomics). In brief, 0.5 ml blood was incubated with 5 ml ACK Lysing Buffer (cat. A1049201, Thermo Scientific) for 5 min at room temperature, followed by adding cold PBS to stop lysis and wash cells. Isolated cells were pelleted and blocked with anti-mouse CD16/CD32 monoclonal antibody (1:100, cat. 14-0161-82, Thermo Scientific) in a 50 µl volume for 10 min at 4 °C to proceed to hashtag staining.

For the single-cell suspensions of the brain, each hemisphere was dissociated by enzymatic digestion with Adult Brain Dissociation Kit (cat. 130-107-677, Miltenyi Biotec) following the manufacturer’s instructions without the step of red blood cell removal. Each hemisphere was cut into small pieces then triturated with a dissociation solution using a GentleMACS dissociator (Miltenyi Biotec). The cell suspension was filtered through a 70 µm filter and washed with DPBS followed by debris removal by gradient centrifugation. For a better cell sorting, the cell suspension was processed to further remove myelin debris with Myelin Removal Beads II (cat. 130-096-733, Miltenyi Biotec) following the manufacturer’s instructions. In brief, cell suspension was incubated with beads then loaded onto a LS Column (cat. 130-042-401, Miltenyi Biotec), which is placed in the magnetic field of a MACS Separator (QuadroMACS Separator, cat. 130-091-051, Miltenyi Biotec). The magnetically labeled myelin was retained within the column and the unlabeled cells ran through. Next, the unlabeled cells were washed and resuspended in 50 µl blocking solution for 10 min at 4 °C to block Fc receptors.

### Cell sorting

To perform fluorescence-activated cell sorting (FACS), brain single-cell suspensions were stained with CD45-PE, CD11b-PE-Cy7, CX3CR1-APC-Cy7, Ly-6C-PerCP-Cy5.5, Ly-6G-FITC. CD45hi cells and microglia (CD45intCD11b+CX3CR1+) were sorted on BD FACS Aria IIIu Cell Sorter (FLI flow cytometry facility, SFig. 1a). Flow cytometry data were analyzed with FlowJo (v10.10.0).

### Single cell RNA sequencing

Isolated blood leukocytes and sorted brain cells from one endpoint were pooled together into one sample to perform 10xGenomics scRNA-seq. To distinguish the origin of cells, blood- and brain-derived cells from different mice were stained with distinct TotalSeq-B hashtag antibodies (0.1µg, BioLegned) in a 50 µl volume for 30 min at 4 °C. Next, cells were washed and resuspended for scRNA-seq.

scRNA-seq was performed at the FLI CF Next Generation Sequencing. scRNA-seq and Cell Surface Protein libraries were generated using the 10x Genomics Chromium X and the Chromium Single Cell 3’ Reagent Kit with 3’ Feature Barcode Kit, Dual Index Kit TT Set A and Dual Index Kit NT Set A, following the manufacturer’s protocols. Single-cell suspensions with oligo-conjugated antibodies were processed in 3 independent batches of 2 samples. Each batch comprised one sample from blood and one from brain, targeting 16,000 cells for blood and 20,000 cells for brain. cDNA yield was assessed on Agilent 4200 TapeStation System with High Sensitivity D5000 ScreenTape (Agilent Technologies, USA) and cDNA was used for 3’ Gene Expression library preparation and Cell Surface Protein library preparation. Final libraries were evaluated using D5000 ScreenTape. The libraries were sequenced on Illumina NovaSeq 6000 System (paired end; R1: 28 bp, R2: 90 bp, I1 and I2: 10 bp). Initial analysis with Cellranger (version: 7.2.0) using bcl2fastq v2.20.0.422 and Mouse reference, GRCm39 (GENCODE v33/Ensembl 110 annotations) estimated 1,500-19,600 cells per sample.

### scRNA-seq data pre-processing

Demultiplexed FASTQ files of brain and blood of three timepoints (day 1, 7, 14) were used as input of *CellRanger* (v7.2) ^108^ to build a genes × cells count matrix separately. Then, we created Seurat Objects using the R package *Seurat* (v5.1.0) ^109^ and performed quality control. We used *HTODemux* function from Seurat to demultiplex mouse samples based on data from cell hashtags. We also used the R package *scDblFinder* (v1.18.0) ^110^ to detect doublets. Cells tagged with a “Singlet” call were kept to QC. Brain cells with fewer than 500 or more than 20000 UMIs, fewer than 200 or more than 6000 genes, more than 5% mitochondrial genes and more than 3% hemoglobin genes were excluded. Blood cells with fewer than 500 or more than 10000 UMIs, fewer than 200 or more than 4000 genes, more than 5% mitochondrial genes and more than 5% hemoglobin genes were excluded. Finally, each time point of the brain samples had an individual Seurat object, and the same applied to blood samples.

### scRNA-seq data analysis

Counts of each Seurat object were normalized using the *SCTransform* function with removing confounding sources of mitochondrial mapping percentage. For each tissue dataset (brain and blood), Seurat objects from three time points were merged into a single Seurat object using Seurat’s *merge* function, followed by principal component analysis (PCA) using *RunPCA* function. Integration and batch effect correction across time points were then performed using *Harmony* (v1.2.3) ^111^, applied to the merged object with the first 30 principal components for the brain dataset and the first 20 principal components for the blood dataset. Next, the neighbor graph of the integrated object was constructed using the *FindNeighbors* function call on harmony reduction on the same principal components, then was used to perform graph-based clustering by *FindClusters* function with a resolution setting of 1 for the brain and 0.9 for the blood. UMAP was calculated on the same principal components using the *RunUMAP* function and was visualized in a two-dimensional space by the *DimPlot* function.

Cell type annotation was initially performed using the *SingleR* package (v2.2.0) ^112^ with four reference datasets (ImmGen ^113^, BrainImmuneAtlas ^114^, TabulaMuris ^115^, and StrokeVis ^6^). Annotations were further validated by integrating cell-type–specific signature scores calculated with the *AddModuleScore* function in Seurat and the expression of marker genes (see STab. 1). Based on these, we identified microglia, monocytes, macrophages, dendritic cells (DCs), neutrophils, T cells, B cells, NK cells, and Innate lymphoid cells (ILCs) for the brain dataset, and neutrophils, B cells, T cells, NK cells, monocytes, DCs, and basophils for the blood dataset. Subpopulation identities of DCs and T cells were assigned based on canonical marker combinations curated from published literature (see STab. 1).

To further characterize stroke-associated transcriptional changes within individual cell types, we generated cell type–specific count matrices and repeated normalization, principal component analysis, *FindNeighbors* using the first 20 principal components, and *FindClusters with* a resolution setting of 0.5. Cluster-specific marker genes were identified using the *FindAllMarkers* function in Seurat. To prioritize robust marker genes, we applied a stringent filtering strategy that considered expression prevalence, effect size, and statistical significance. Differentially expressed genes (DEGs) were retained only if they were detected in at least 75% of cells in the target population (pct ≥ 0.75), showed a minimum average expression difference of 1.5-fold (avg_log2FC ≥ log2(1.5)), and remained significant after Bonferroni correction (p_val_adj < 0.05), and were ranked by avg_log2FC.

In addition, within each cell type, differential gene expression analysis between MCAo and Sham conditions was performed independently at each time point using the *FindMarkers* function, with experimental conditions used as the grouping variable. The DEGs detected in at least 10% of cells in the target population (pct ≥ 0.1), showed a minimum average expression difference of 1.5-fold (avg_log2FC ≥ log2(1.5)), and remained significant after Bonferroni correction (p_val_adj < 0.05) were identified as condition-associated DEGs. Z-scores were calculated based on the average gene expression across clusters or experimental conditions and displayed in heatmaps.

### Gene Ontology (GO) term enrichment analysis

DEGs identified between MCAo and sham microglia at each time point were used for GO term enrichment analysis. Genes with detection in at least 10% of cells in either group (pct.1 ≥ 0.1 or pct.2 ≥ 0.1) were retained as the background candidate space for downstream enrichment analysis. Within each time point, genes with avg_log2FC ≥ log2(1.5) and adjusted P < 0.05 were classified as upregulated, whereas genes with avg_log2FC ≤ -log2(1.5) and adjusted P < 0.05 were classified as downregulated; all remaining genes were classified as non-significant changed genes. Time-point-specific results were merged into a DEG scale matrix across all time points.

Analysis was performed separately for upregulated and downregulated gene sets using the *compareCluster* function in the *clusterProfiler* package (v4.14.6) ^116^ with the formula interface *(Gene ∼ Timepoint)* and *enrichGO* as the enrichment function. Mouse gene annotation was obtained from *org.Mm.eg.db* (v3.19.1) ^117^. The background universe was defined as the genes that passed the minimum expression detection filter (pct.1 ≥ 0.1 or pct.2 ≥ 0.1) in the corresponding MCAo-versus-sham comparisons across the three time points. GO enrichment was conducted independently for Biological Process (BP), Molecular Function (MF), and Cellular Component (CC) categories. Terms were considered enriched at pvalueCutoff < 0.05 and qvalueCutoff < 0.2. Unless otherwise specified, default parameters in clusterProfiler were used, including Benjamini–Hochberg correction for multiple testing.

### Cell trajectory inference

We used the *dyno* R package (v0.1.2) ^118^ to conduct the trajectory inference analysis for microglia. Gene expression values in the RNA assay were log-normalized, and 3000 highly variable genes were identified, followed by data scaling, PCA, nearest-neighbor graph construction, and UMAP embedding. The normalized expression matrix and raw count matrix from the RNA assay were exported and wrapped into a dyno dataset using *wrap_expression* function, and prior information was provided by specifying the starting cells and PCA coordinates. All cells from the day1_sham group were defined as the starting population. Based on dyno guideline recommendations, *Slingshot* ^119^ was selected as the trajectory inference method and was run through the dyno framework. Trajectories were visualized in two-dimensional UMAP space.

### Cell communication analysis

Brain and peripheral blood datasets were merged and were partitioned into subsets corresponding to each annotated cell type. We inferred cell-cell communication networks using *CellChat* R package (v2.2.0) ^120^. The built-in mouse ligand–receptor database was used after excluding the non-protein signaling category. The processed Seurat object was split by time point and condition, and CellChat analysis was performed separately for each subset using the RNA assay. Overexpressed genes and interactions were identified using *identifyOverExpressedGenes* and *identifyOverExpressedInteractions*, respectively. Communication probabilities were inferred using *computeCommunProb* with the triMean method, and interactions involving cell groups with fewer than 15 cells were removed using *filterCommunication*. Pathway-level communication probabilities were calculated with *computeCommunProbPathway*, and aggregated communication networks were generated with *aggregateNet*. The inferred networks were visualized as circle plots showing the number or strength of interactions by using *netVisual_diffInteraction*.

### Single cell pathway enrichment analysis

To characterize immune-related functional programs across major immune cell types, single-sample gene set enrichment analysis (ssGSEA) was performed on the integrated Seurat objects of monocytes, B cells, neutrophils, T cells, and NK cells using the *escape* R package (v2.6.2) ^121^ together with the *GSVA* (v1.52.3) ^122^ framework. For each cell type, the RNA assay was used as input, and enrichment scores were calculated at single-cell resolution with *runEscape* function using the *ssGSEA* method. Three curated GO-slim–based gene sets were evaluated: inflammatory response (GO:0006954; 661 genes), cytokine-mediated signaling pathway (GO:0019221; 439 genes), and leukocyte migration (GO:0050900; 252 genes). Enrichment scores were normalized using the built-in normalization option in *escape*. Mean pathway scores were then summarized by group and visualized as heatmaps, with row-wise Z-score scaling.

### Brain-blood cell distance analysis

A custom per-mouse kNN distance analysis was performed to quantify the transcriptomic similarity between brain and paired blood cells. For each mouse, brain and blood cells of the same annotated cell type were subsetted and processed independently in Seurat using log-normalization, variable feature selection, scaling, and PCA. Euclidean distances in PCA space were computed with RANN::nn2(v2.6.2) ^123^using the first 20 PCs. For each brain cell, the mean distance to its five nearest blood-cell neighbors from the same mouse was calculated (brain-to-blood distance). As a within-mouse baseline, blood-to-blood distances were calculated analogously among blood cells after excluding self-hits. A robust z-score was additionally calculated using the median and median absolute deviation (MAD) of the blood-to-blood distances. Lower z-scores indicate greater transcriptomic similarity of brain cells to the paired blood reference population.

### Sample preparation for proteomics analysis

Tissues were homogenized in PBS containing protease and phosphoatase inhibitors (cOmplete™ ULTRA Tablets and PhosSTOP™, Roche) using a bead beater (Precellys ®, Bertin Technologies, France). Afterwards a lysis buffer (fc 4% SDS, 100 mM HEPES, pH 8.5, 50 mM DTT) was added to 20 μg of each sample. Samples were then boiled at 95°C for 10 min and sonicated using a tweeter. Reduction was followed by alkylation with 200 mM iodoacetamide (IAA, final concentration 15 mM) for 30 min at room temperature in the dark. Samples were acidified with phosphoric acid (final concentration 2.5%), and seven times the sample volume of S-trap binding buffer was added (100 mM TEAB, 90% methanol). Samples were bound either on a 96-well S-trap mini plate (Protifi) and washed three times with a binding buffer. Trypsin in 50 mM TEAB pH 8.5 was added to the samples (1 µg per sample) and incubated for 1 h at 47°C. The samples were eluted in three steps with 50 mM TEAB pH 8.5, elution buffer 1 (0.2% formic acid in water) and elution buffer 2 (50% acetonitrile and 0.2% formic acid). The eluates were dried using a speed vacuum centrifuge (Eppendorf Concentrator Plus, Eppendorf AG, Germany) and stored at -20° C. Before analysis, samples were reconstituted in in MS Buffer (5% acetonitrile, 95% Milli-Q water, with 0.1% formic acid), spiked with iRT peptides (Biognosys, Switzerland) and loaded on Evotips (Evosep) according to the manufacturer’s instructions. In short, Evotips were first washed with Evosep buffer B (acetonitrile, 0.1% formic acid), conditioned with 100% isopropanol and equilibrated with Evosep buffer A. Afterwards, the samples were loaded on the Evotips and washed with Evosep buffer A. The loaded Evotips were topped up with buffer A and stored until the measurement.

### LC-MS Data independent analysis (DIA)

Peptides were separated using the Evosep One system (Evosep, Odense, Denmark) equipped with a 15 cm x 150 μm i.d. packed with a 1.5 μm Reprosil-Pur C18 bead column (Evosep performance, EV-1137, Denmark). The samples were run with a pre-programmed proprietary Evosep gradient of 44 min (30 samples per day) using water and 0.1% formic acid and solvent B acetonitrile and 0.1% formic acid as solvents. The LC was coupled to an Orbitrap Exploris 480 (Thermo Fisher Scientific, Bremen, Germany) using PepSep Sprayers and a Proxeon nanospray source. The peptides were introduced into the mass spectrometer via a PepSep Emitter 360-μm outer diameter × 20-μm inner diameter, heated at 300°C, and a spray voltage of 2 kV was applied. The injection capillary temperature was set at 300°C. The radio frequency ion funnel was set to 30%. For DIA data acquisition, full scan mass spectrometry (MS) spectra with a mass range of 350–1650 m/z were acquired in profile mode in the Orbitrap with a resolution of 120,000 FWHM. The default charge state was set to 2+, and the filling time was set at a maximum of 20 ms with a limitation of 3 × 106 ions. DIA scans were acquired with 40 mass window segments of differing widths across the MS1 mass range. Higher collisional dissociation fragmentation (normalized collision energy 30%) was applied, and MS/MS spectra were acquired with a resolution of 30,000 FWHM with a fixed first mass of 200 m/z after accumulation of 1 × 106 ions or after filling time of 45 ms (whichever occurred first). Data was acquired in profile mode. For data acquisition and processing of the raw data, Xcalibur 4.4 (Thermo) and Tune version 4.0 were used.

### Proteomic data processing

DIA raw data were analyzed using the directDIA pipeline in Spectronaut v.20 (Biognosys, Switzerland) with BGS settings besides the following parameters: Protein LFQ method= QUANT 2.0, Proteotypicity Filter = Only protein group specific, Major Group Quantity = Median peptide quantity, Minor Group Quantity = Median precursor quantity, Data Filtering = Qvalue, Normalizing strategy = Local Normalization. The data were searched against a species specific (Mus musculus, 16,747 entries, v. 160106) and a contaminants (247 entries) Swissprot database. The identifications were filtered to satisfy FDR of 1 % on peptide and protein level. The data were then exported and further analyzed in Rstudio using MSStats ^124^ removing sparse precursors and with a cutoff for missing values of 0.5 for quantification of proteins

### Proteomics analysis

All proteomics analyses were performed in R (version 4.5.2) using tidyverse (v.2.0.0) ^125^, ComplexHeatmap (v.2.26.1) ^126^, AnnotationDbi (v.1.72.0) ^127^ and further packages described below. Log2-transformed protein-by-sample intensity matrices were obtained for five intestinal segments (duodenum, jejunum, terminal ileum, caecum and proximal colon) and mesenteric lymph nodes (MLN) from 34 animals across three post-operative timepoints (day 1, 7 and 14), yielding 176 samples (25 to 34 per tissue). Proteins with missing values in more than 50% of samples within each tissue were removed, as were proteins with zero variance. Filtering was performed independently per tissue. As animals were sacrificed across seven dates, we assessed the contribution of sacrifice date to proteome variation. PERMANOVA ^128^ (“vegan::adonis2” ^129^; Euclidean distance on complete-case samples; 999 permutations) on uncorrected data indicated that sacrifice date explained 11–26% of total proteome variance across tissues. To account for this batch structure, we applied surrogate variable analysis (SVA) ^130,131^ using the Buja–Eyuboglu permutation method ^132^, resulting in 2 to 5 surrogate variables per tissue. The full model (∼0 + group, where the group encodes the six condition-by-timepoint combinations) protected both main effects and the condition-by-timepoint interaction from being absorbed into surrogate variables. For differential expression testing, surrogate variables were included as covariates directly in the linear model rather than being regressed out of the expression matrix, to avoid removing biological signal correlated with batch structure. Two additional approaches were used for sensitivity analysis: uncorrected data and data corrected for sacrifice date using removeBatchEffect from limma ^133^ with sacrifice date as a categorical batch variable. UniProt accession identifiers were mapped to mouse gene symbols using three complementary sources: the UniProt REST API ^134^ (primary accession to gene name), the org.Mm.eg.db ^117^ Bioconductor annotation package (which additionally resolves secondary and obsolete accessions), and manual curation for 20 entries including immunoglobulin constant/variable region chains and proteins missed by both automated sources. Non-mouse proteins (31 contaminants including human keratins, bovine serum albumin and porcine trypsin) were identified via organism annotation from the UniProt API and excluded. The union of all three sources yielded gene symbol assignments for 7,344 of 7,382 unique UniProt IDs (99.5%). The remaining 38 unmapped entries comprised the 31 non-mouse contaminants and 7 demerged or obsolete identifiers. For identifiers mapping to multiple gene symbols (172 IDs), all matching symbols were retained. When multiple UniProt identifiers mapped to the same gene symbol, per-sample median intensity was taken across the corresponding protein entries, producing gene-level expression matrices per tissue.

Differential expression analysis was performed per tissue using limma on uncorrected, log2-transformed, gene-level expression matrices with surrogate variables included as covariates in the design matrix. For each tissue, a linear model with condition-by-timepoint group as the explanatory variable (∼0 + group, six levels) and tissue-specific surrogate variables was fitted, and three contrasts comparing MCAo versus sham at each timepoint (d1, d7, d14) were extracted. P-values were adjusted using the Benjamini–Hochberg method ^135^. To estimate the proportion of genes with true signal regardless of individual significance, π₀ was computed using the qvalue R package ^136^ on raw p-values for each tissue, timepoint and correction method. The complement (1 − π₀) estimates the fraction of genes with non-null effects.

In a hypothesis-driven approach, we tested a series of pre-defined biological hypotheses using curated gene sets from Gene Ontology Biological Process (GO:BP) ^137^ terms obtained via msigdbr ^138^. The following gene sets were evaluated:

- tight junction organization (GOBP_TIGHT_JUNCTION_ORGANIZATION, 87 genes),

- extracellular matrix organization (GO:0030198 via org.Mm.eg.db, 337 genes),

- PPAR signaling

(GOBP_PEROXISOME_PROLIFERATOR_ACTIVATED_RECEPTOR_SIGNALING_ PATHWAY, 28 genes),

- pyruvate metabolism (GOBP_PYRUVATE_METABOLIC_PROCESS, 126 genes),

- oxidative phosphorylation (GOBP_OXIDATIVE_PHOSPHORYLATION, 143 genes),

- fatty acid β-oxidation (GOBP_FATTY_ACID_BETA_OXIDATION, 72 genes),

- epithelial cell proliferation (GOBP_EPITHELIAL_CELL_PROLIFERATION, 449 genes),

- wound healing (GOBP_WOUND_HEALING, 438 genes),

- leukocyte migration (GOBP_LEUKOCYTE_MIGRATION, 381 genes),

- myeloid leukocyte migration (GOBP_MYELOID_LEUKOCYTE_MIGRATION, 221 genes),

- Toll-like receptor signaling pathway (GOBP_TOLL_LIKE_RECEPTOR_SIGNALING_PATHWAY, 79 genes),

- regulation of isotype switching (GOBP_REGULATION_OF_ISOTYPE_SWITCHING, 38 genes),

- MHC class I antigen presentation (GOBP_ANTIGEN_PROCESSING_AND_PRESENTATION_OF_PEPTIDE_ANTIGE N_VIA_MHC_CLASS_I, 35 genes),

- MHC class II antigen presentation (GOBP_ANTIGEN_PROCESSING_AND_PRESENTATION_OF_EXOGENOUS_PE PTIDE_ANTIGEN_VIA_MHC_CLASS_II, 23 genes),

- T cell activation (GOBP_T_CELL_ACTIVATION, 574 genes),

- Th17 cell differentiation (GOBP_T_HELPER_17_CELL_DIFFERENTIATION, 40 genes),

- ãä T cell activation (GOBP_GAMMA_DELTA_T_CELL_ACTIVATION, 26 genes),

- IL-17 production (GOBP_INTERLEUKIN_17_PRODUCTION, 41 genes).

In addition, the Reactome ^139^ pathway for PGC-1α activation (REACTOME_ACTIVATION_OF_PPARGC1A_PGC_1ALPHA_BY_PHOSPHORYLATION, 10 genes) was included as a more specific test of the colonocyte metabolic switch hypothesis. Note that the GO:BP PPAR signaling term encompasses all PPAR isoforms (alpha, beta/delta, gamma) and is not specific to PPAR-gamma. For each gene set, tissue and timepoint, a per-sample summary score was computed as the mean z-score across all detected genes in the set (z-scored within tissue across samples). MCAo versus sham scores were compared using two-sided Welch’s t-tests ^140^. Analyses were performed on both SVA-corrected (primary) and uncorrected (sensitivity) data (STab. 6).

Gene set enrichment analysis (GSEA) ^95^ was performed using the fgsea R package ^141^ with 186 KEGG ^142^ pathways from the MSigDB database (human gene sets mapped to mouse orthologs via msigdbr). For each tissue and timepoint, proteins were ranked by the SVA-adjusted moderated t-statistic from the limma analysis described above and GSEA was run with 10,000 permutations (minSize = 15, maxSize = 500). P-values were adjusted using BH correction. After identifying pathways significant in at least one tissue-by-timepoint combination, we applied pathway collapsing (collapsePathways from fgsea) to remove redundant gene sets sharing leading edge genes. We then applied an entry-level direction-concordance filter: each tissue × timepoint × pathway entry was retained only where the sign of enrichment in the SVA-corrected analysis matched the parallel uncorrected analysis; reversed entries were masked from the heatmap. 364 of 1,242 entries (29%) were masked; 69 of 72 collapsed pathways retained at least one direction-concordant entry and are shown. Clustering of pathways was performed on the full unmasked SVA NES matrix, so masking affects display only.

### Analysis of the gut microbiota

Stool samples were collected from n=4 sham and n=6 MCAo animals (days: d−1, d1, d3, d7, d14), where d−1 represents the pre-surgical baseline. Metatranscriptomic sequencing was performed on samples from all 5 timepoints, yielding a total of 46 libraries. Metagenomic sequencing was performed on day 14 samples from all animals, yielding 10 libraries.

### Metagenomics data analysis

Sequencing reads were processed using the ATLAS metagenome assembly pipeline (v2.19.0, default settings) ^143^. Raw reads were quality control to remove adapter sequences, PCR duplicates, and contaminations. Quality-controlled reads were assembled de novo using both SPAdes (v4.0) ^144^, and the resulting contigs were further filtered by mapping the quality-controlled reads back to the assemblies to retain only high-confidence contigs. Genome binning was performed using VAMB (v3.0.9) ^145^, and the resulting bins were consolidated using DAS Tool to generate non-redundant metagenome-assembled genomes (MAGs) per sample. Bin quality was assessed with CheckM2 (v1.0.2) ^146^, and MAGs were retained based on completeness and contamination thresholds consistent with established standards (completeness ≥50%, contamination ≤10% for medium-quality; completeness ≥90%, contamination ≤5% for high-quality MAGs). Protein-coding genes were predicted on the filtered contigs using Prodigal, and taxonomic classification of MAGs was performed using the Genome Taxonomy Database Toolkit (GTDB-Tk v2.4) ^147^.

### Prediction of facultative anaerobes

Facultative anaerobes were predicted using Bacdive AI (github release a1bef3e from 9/2024) ^148^. If neither the anaerobic nor the aerobic trait could be predicted, we considered the organisms to be facultative, as suggested by the BacDive AI publication. We aggregate the abundance of bacteria that were predicted to be facultative anaerobes and tested their influence in a linear mixed effect model with time and condition as fixed and cage and replicate as random factors. We then checked the impact of fixed factors using ANOVA with Satterthwaite’s method of the lmerTest (v3.2.1) ^149^ and lme4 (v2.0.1) ^150^ R packages.

### Metatranscriptomics data analysis

An initial quality assessment of raw sequencing reads was performed using FastQC. Adapter trimming and low-quality read filtering were carried out using Trimmomatic (v0.38) ^151^. Host-derived reads were subsequently removed by aligning the trimmed reads to the mouse reference genome (downloaded from NCBI) using Bowtie2 (v2.2.6) ^152^, and all reads mapping to the host genome were discarded. A second FastQC assessment was performed on the processed reads to confirm adequate quality prior to downstream analysis. Ribosomal RNA reads were identified and removed using SortMeRNA (v2.1) ^153^, retaining only putative mRNA reads for subsequent analyses.

### Taxonomic classification and community diversity analysis

A custom reference database was constructed by combining metagenome-assembled genomes (MAGs) derived from the paired metagenomic sequencing data with 55 additional reference genomes retrieved from NCBI. The latter were selected based on high read-assignment rates observed during an initial taxonomic classification performed with Kraken2 and Bracken ^154^, ensuring that the reference database captured the dominant taxa present in the dataset. Quality-controlled metatranscriptomic reads were aligned to this in-house reference database using Bowtie2, and the resulting sequence alignment files were processed with SAMtools. Per-taxon read counts were extracted using the SAMtools idxstats command and normalised by library size to generate a taxon-by-sample abundance matrix. Microbial community diversity was assessed using the vegan package in R. Alpha diversity was quantified using standard indices, and beta diversity was evaluated using both Aitchison and Bray-Curtis dissimilarity metrics to capture compositional and abundance-based differences between communities, respectively. Differential abundance analysis was performed using MaAsLin2 ^155^.

### Gene expression and eifferential expression analysis

Read counts for annotated genomic features were quantified using the featureCounts function from the Subread package ^156^, generating a gene-by-sample count matrix. Differential expression analysis was performed using the dream function within the variancePartition package ^80^, which extends the limma/voom framework ^157^ by fitting linear mixed models to appropriately account for repeated measures and variation attributable to cage effects. This was modelled using a linear mixed model with cell-mean fixed effects for each Condition × timepoint combination (∼ 0 + Condition:time), and random intercepts for biological sample (1|sample) and housing cage (1|Cage) to account for repeated measures/within-subject correlation and cage-level clustering respectively. Hypothesis testing for differences between two or more model coefficients was performed by specifying contrasts evaluated at the time of model fitting, which allowed uncertainty to be carried through consistently across all comparisons. Biologically relevant contrasts were then specified using makeContrasts function and grouped into three categories: (i) within-condition temporal contrasts comparing each timepoint to baseline (day -1) separately in stroke and sham groups, (ii) cross-sectional contrasts comparing stroke to sham at each timepoint, and (iii) interaction contrasts formally testing whether the change from baseline differed between stroke and sham at each timepoint. Empirical Bayes moderation was applied using eBayes with robust = TRUE to stabilise variance estimates. Multiple testing correction was performed using the Benjamini-Hochberg false discovery rate method, and genes with an adjusted p-value ≤ 0.05 and an absolute log2 fold change ≥ 1 were considered differentially expressed.

### Functional enrichment analysis

To assess the biological relevance of differentially expressed genes, over-representation analysis was performed against Gene Ontology (GO) and MetaCyc pathways ^158^. To further capture transcriptional changes beyond individual gene-level significance thresholds, gene set enrichment analysis (GSEA) was conducted using *fgsea* on a full ranked gene list derived from the differential expression analysis. For each contrast, genes were ranked by the standardized z-score (z.std) derived from the dream mixed model. This metric transforms the gene-specific moderated t-statistic to the standard normal scale using each gene’s Satterthwaite degrees of freedom, making the ranking comparable across genes regardless of their individual uncertainty. The enrichment scores were computed using 10000 (nPermSimple) permutations, with gene set limits as 10 to 500 genes. To improve biological interpretability and identify broad pathways with distinct core genes, *collapsPathways* function was used on significant pathways (padj < 0.05) obtained from *fgsea*. For better interpretation, MetaCyc Level 1 and Level 2 hierarchies were then explored and used to summarize the count of significant pathways per level.

## Data availability

Currently, all data and analysis scripts are available from the corresponding author upon reasonable request.

## Acknowledgements

We thank the Core Facility Next Generation Sequencing (Ivonne Görlich and Martin Bens) for their support in library preparation and sequencing, CF Proteomics, CF Flow Cytometry (Kathrin Schubert, Johanna Schleep and Michael Müller) and the Mouse Facility at the Leibniz Institute on Aging, Fritz Lipmann Institute (FLI) for supporting our experiments. HMD is funded by Carl-Zeiss-Stiftung (P2021-00-007). KW and HMD received a conference grant from the Fritz Thyssen Foundation for the organization of the Jena Microbiome Meeting (2023) that sparked relevant collaborations. KW, HMD and JZ are associated with the Cluster of Excellence - Balance of the Microverse and BK was supported by the Cluster of Excellence - Balance of the Microverse (project grant to HMD). This work was funded by the Deutsche Forschungsgemeinschaft (DFG, German Research Foundation) under Germany’s Excellence Strategy – EXC 2051 – Project-ID 390713860 and supported by the ERA-NET NEURON network (DFG, MOODYGUT project #561967932 to KW).

