## Supplementary Figures for "Temporal multi-omic profiling of immune, gut, and microbiome responses to ischemic stroke reveals convergence of host and microbial perturbations one week after brain injury"

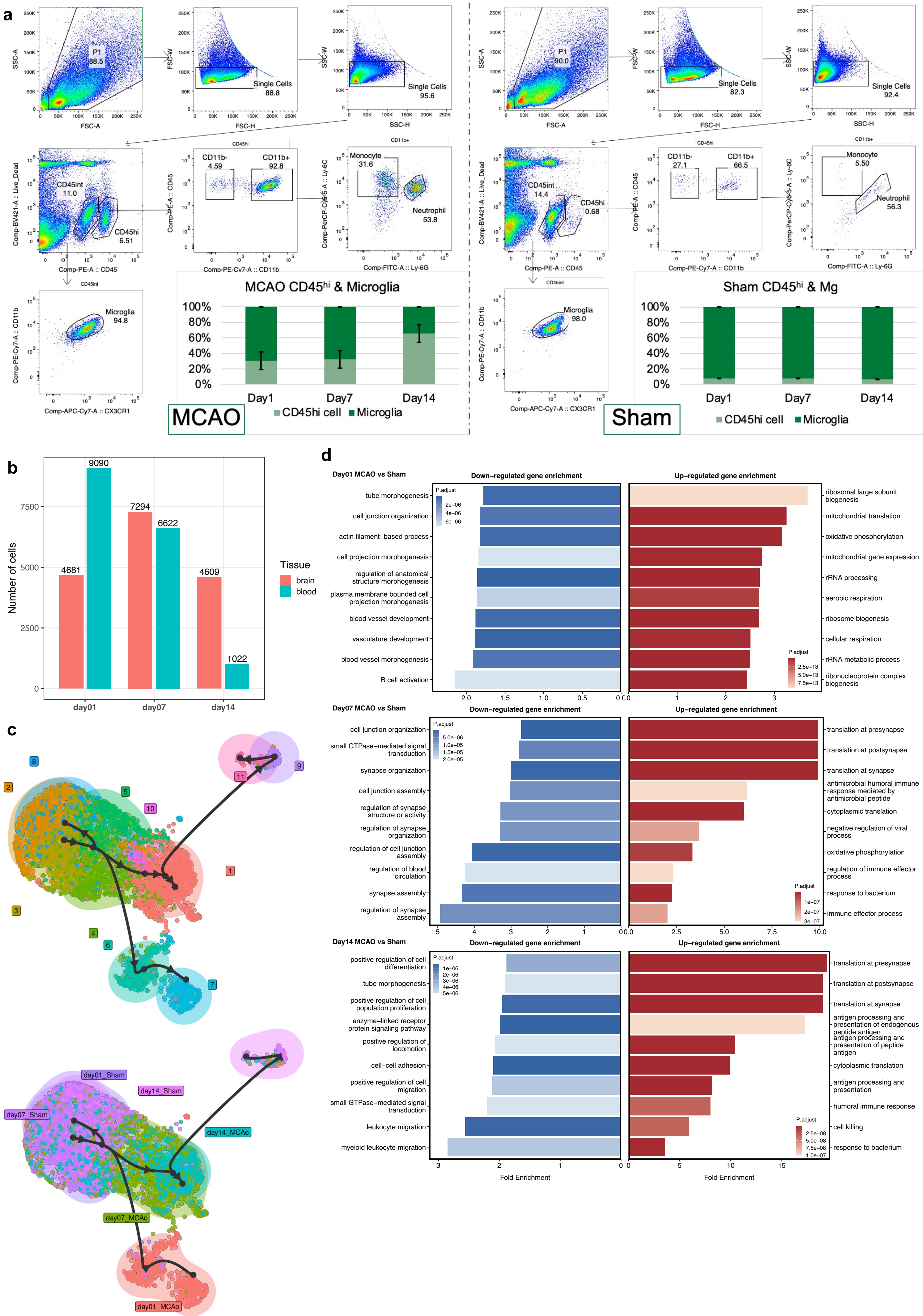
a
P1 88.5
Single Cells 88.8
Single Cells 95.6
SSC-A
FSC-A
FSC-W
FSC-H
SSC-W
SSC-H
CD45int 11.0
CD45hi 6.51
Comp-BV421-A :: Live_Dead
Comp-PE-A :: CD45
CD45hi
CD11b- 4.59
CD11b+ 92.8
Comp-PE-Cy7-A :: CD11b
CD11b+
Monocyte 31.6
Neutrophil 53.8
Comp-PerCP-Cy5-5-A :: Ly-6C
Comp-FITC-A :: Ly-6G
CD45int
Microglia 94.8
Comp-APC-Cy7-A :: CX3CR1
MCAO CD45hi & Microglia
100%
80%
60%
40%
20%
0%
Day1
Day7
Day14
CD45hi cell
Microglia
MCAO
P1 90.0
Single Cells 82.3
Single Cells 92.4
CD45int 14.4
CD45hi 0.68
CD11b- 27.1
CD11b+ 66.5
Monocyte 5.50
Neutrophil 56.3
Microglia 98.0
Sham CD45hi & Mg
Sham
b
Number of cells
4681
9090
7294
6622
4609
1022
day01
day07
day14
Tissue
brain
blood
c
1
2
3
4
5
6
7
8
9
10
11
day01_Sham
day14_Sham
day07_Sham
day14_MCAO
day07_MCAO
day01_MCAO
d
Day01 MCAO vs Sham
Down−regulated gene enrichment
Up−regulated gene enrichment
tube morphogenesis
cell junction organization
actin filament−based process
cell projection morphogenesis
regulation of anatomical structure morphogenesis
plasma membrane bounded cell projection morphogenesis
blood vessel development
vasculature development
blood vessel morphogenesis
B cell activation
P.adjust
2e−06
4e−06
6e−06
ribosomal large subunit biogenesis
mitochondrial translation
oxidative phosphorylation
mitochondrial gene expression
rRNA processing
aerobic respiration
ribosome biogenesis
cellular respiration
rRNA metabolic process
ribonucleoprotein complex biogenesis
2.5e−13
5.0e−13
7.5e−13
Day07 MCAO vs Sham
cell junction organization
small GTPase−mediated signal transduction
synapse organization
cell junction assembly
regulation of synapse structure or activity
regulation of synapse organization
regulation of cell junction assembly
regulation of blood circulation
synapse assembly
regulation of synapse assembly
5.0e−06
1.0e−05
1.5e−05
2.0e−05
translation at presynapse
translation at postsynapse
translation at synapse
antimicrobial humoral immune response mediated by antimicrobial peptide
cytoplasmic translation
negative regulation of viral process
oxidative phosphorylation
regulation of immune effector process
response to bacterium
immune effector process
1e−07
2e−07
3e−07
Day14 MCAO vs Sham
positive regulation of cell differentiation
tube morphogenesis
positive regulation of cell population proliferation
enzyme−linked receptor protein signaling pathway
positive regulation of locomotion
cell−cell adhesion
positive regulation of cell migration
small GTPase−mediated signal transduction
leukocyte migration
myeloid leukocyte migration
1e−06
2e−06
3e−06
4e−06
5e−06
translation at presynapse
translation at postsynapse
translation at synapse
antigen processing and presentation of endogenous peptide antigen
antigen processing and presentation of peptide antigen
cytoplasmic translation
antigen processing and presentation
humoral immune response
cell killing
response to bacterium
2.5e−08
5.0e−08
7.5e−08
1.0e−07
Fold Enrichment
Fold Enrichment

**Supplementary Figure 1. Flow Cytometry strategy and microglia transcriptome profile. (a)** Gating strategy used to identify and sort $CD45^{hi}$ cells and microglia in single cell suspensions of mouse brains from MCAo (*left*) and Sham (*right*) mice. Singlets were gated based on forward and side scatter. Live $CD45^{int}$ and $CD45^{hi}$ cells were gated based on DAPI staining and CD45 expression. Microglia ($CD45^{int}CD11b^{+}CX3CR1^{+}$) were identified in $CD45^{int}$ population based on CD11b and CX3CR1 expression. Analysis of Ly6G and LY6C expression was performed in $CD45^{hi}CD11b^{+}$ population to verify the presence of neutrophils ($Ly6G^{+}$) and monocytes ($Ly6C^{+}Ly6G^{-}$). Bar plots showing relative percentage of $CD45^{hi}$ cells and microglia. **(b)** Bar plot showing number of cells from brain and blood at different time points in our single-cell data after quality control. **(c)** Pseudotime inference of microglial clusters (*top*) or microglia experimental groups (*bottom*) showing a transcriptional progression of microglia from sham-enriched, homeostatic-like states (clusters 2, 3, and 8) toward MCAo-associated activated states. **(d)** GO biological process enrichment analysis of time point-specific microglial DEGs in MCAo versus sham mice. For each time point, GO biological process (GO-BP) over-representation analysis was performed separately for MCAo-enriched genes (up-regulated) and sham-enriched genes (down-regulated) identified from the MCAo-versus-sham comparison in microglia. The background universe was defined as genes tested for differential expression with detection in at least 10% of cells ($pct \geq 0.1$). Input gene sets consisted of DEGs with $pct \geq 0.1$, $|log2FC| \geq log2(1.5)$, and Bonferroni-adjusted $p < 0.05$ in the respective condition. Top 10 enriched GO-BP terms are shown for sham-enriched genes (left) and MCAo-enriched genes (right). Bar length indicates fold enrichment, and bar colour denotes the adjusted p value.

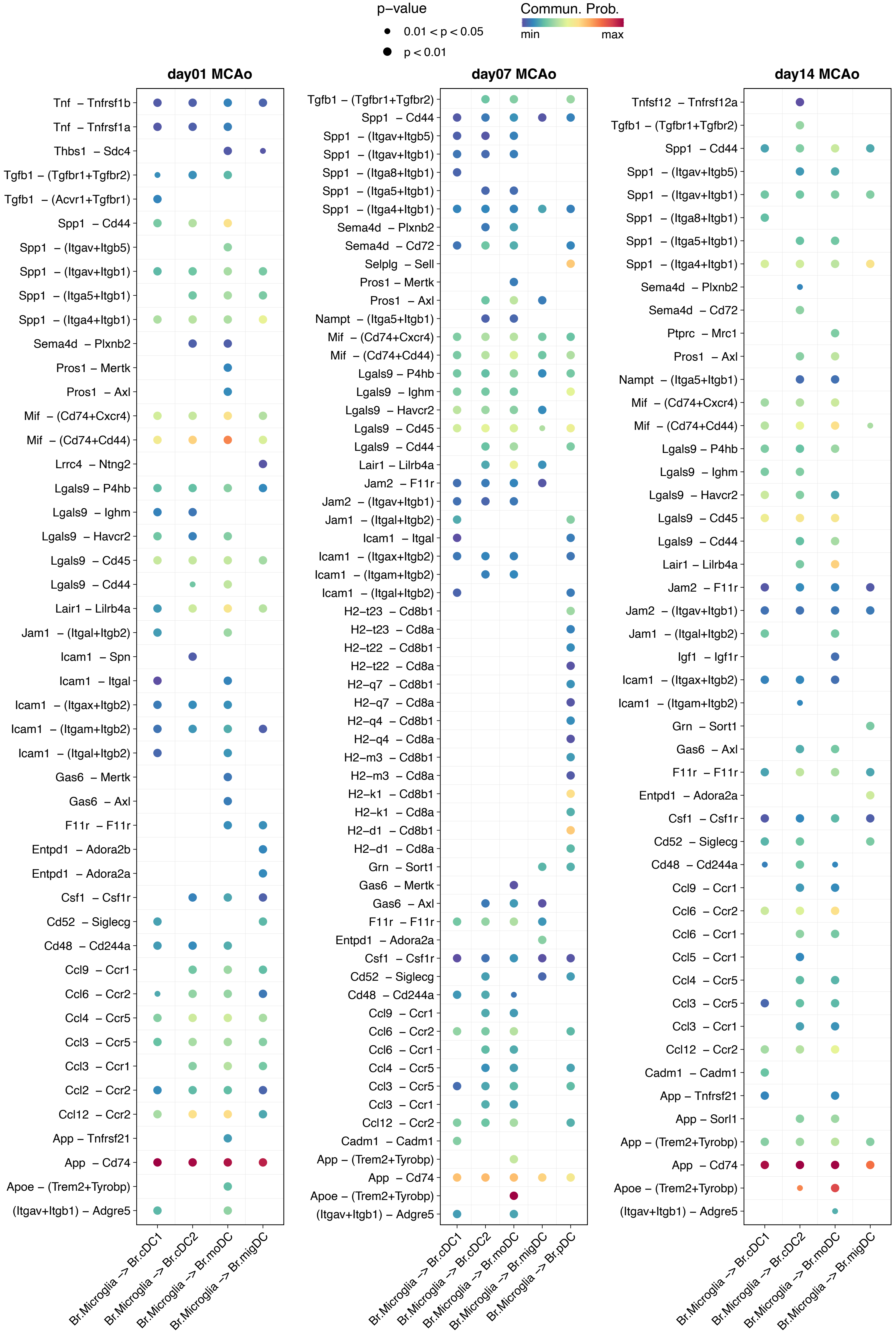

p−value
0.01 < p < 0.05
p < 0.01
Commun. Prob.
min
max
day01 MCAo
day07 MCAo
day14 MCAo
Br.Microglia -> Br.cDC1
Br.Microglia -> Br.cDC2
Br.Microglia -> Br.moDC
Br.Microglia -> Br.migDC
Br.Microglia -> Br.pDC

**Supplementary Figure 2. Microglia - DCs ligand-receptor interactions based on CellChat analysis.** Bubble plots showing ligand-receptor interactions from microglia to DCs in MCAo of day 1 (*left*), day 7 (*middle*) and day 14 (*right*). Rows represent senders and receivers. Columns represent interactions. Bubble size indicates the p-value of interactions. Color intensity indicates the communication probabities (Commun.Prob.) of interactions.

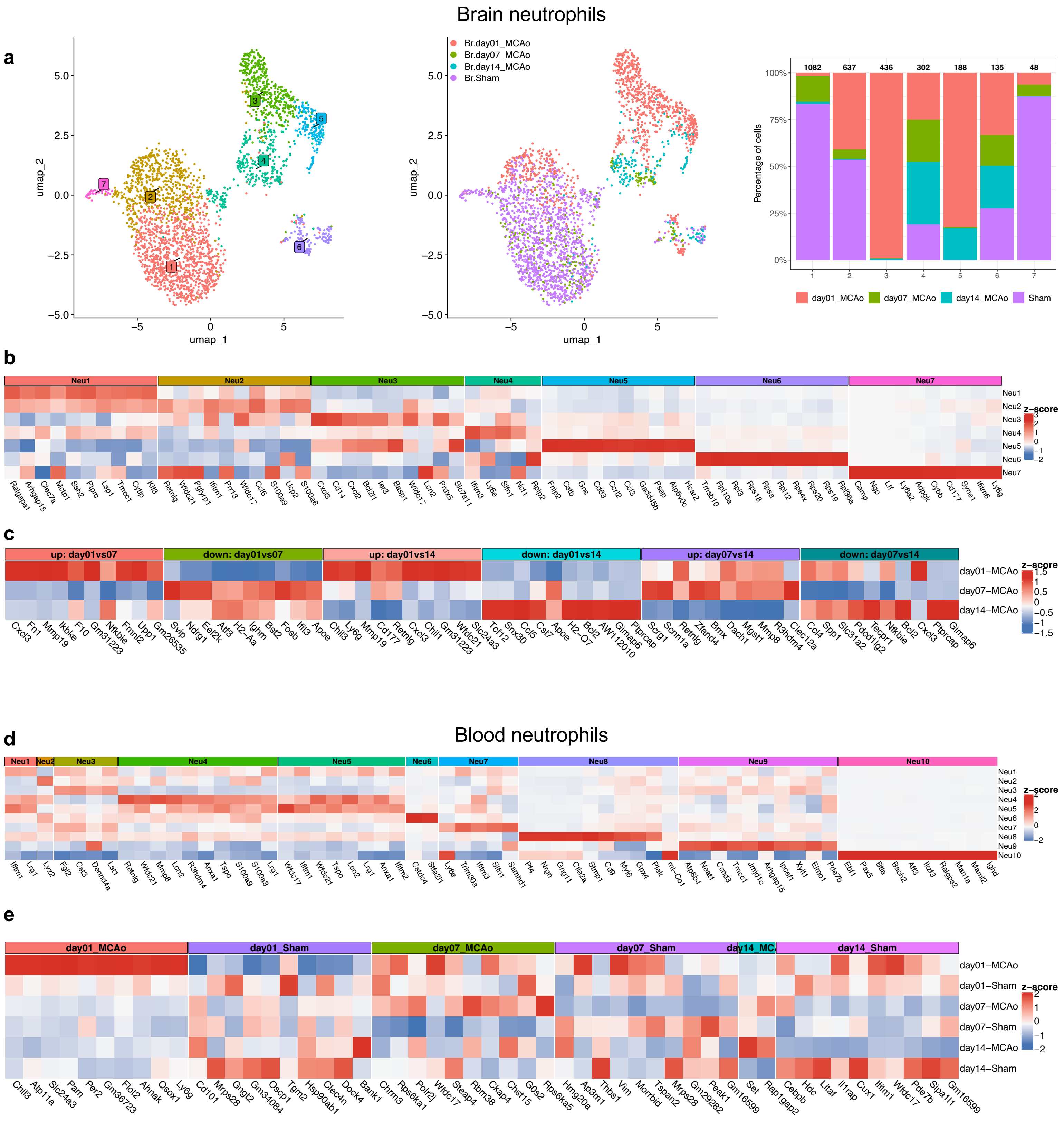

Brain neutrophils
a
umap_2
umap_1
Br.day01_MCAo
Br.day07_MCAo
Br.day14_MCAo
Br.Sham
Percentage of cells
1082 637 436 302 188 135 48
day01_MCAo day07_MCAo day14_MCAo Sham
b
Neu1 Neu2 Neu3 Neu4 Neu5 Neu6 Neu7
z-score
Ralgapa1 Arhgap15 Clec7a Mcpt1 Ssh2 Ptprc Lsp1 Tmcc1 Cyfip Klf3 Retnlg Wfdc21 Pglyrp1 Ifitm1 Prr13 Wfdc17 Ccl6 S100a9 Ucp2 S100a6 Cxcl3 Ccl4 Cxcl2 Bcl2l1 Ier3 Basp1 Wfdc17 Lcn2 Prok2 Slc7a11 Ifitm3 Ly6e Slfn1 Ncf1 Rplp2 Fnip2 Cstb Gns Cd63 Ccrl2 Ccl3 Gadd45b Psap Atp6v0c Hcar2 Tmsb10 Rpl10a Rpl3 Rps18 Rpsa Rpl12 Rps4x Rps20 Rps19 Rpl36a Camp Ngp Ltf Ly6a2 Adpgk Cybb Cd177 Syne1 Ifitm6 Ly6g
c
up: day01vs07 down: day01vs07 up: day01vs14 down: day01vs14 up: day07vs14 down: day07vs14
day01−MCAo day07−MCAo day14−MCAo
Cxcl3 Fn1 Mmp19 Ikbke F10 Gm31223 Nfkbie Fnml2 Upp1 Gm26535 Svip Ndrg1 Eef2k Atf3 H2−Aa Ighm Bst2 Fosb Ifit3 Apoe Chil3 Ly6g Mmp19 Cd177 Retnlg Cxcl3 Chil1 Gm31223 Wfdc21 Slc24a3 Tcf12 Snx30 Ccl5 Cst7 Apoe H2−Q7 Bcl2 AW112010 Gimap6 Ptprcap Scrg1 Scnn1a Retnlg Zfand4 Bmx Dach1 Mgst1 Mmp8 R3hdm4 Clec12a Ccl4 Spp1 Slc31a2 Tecpr1 Pdcd1lg2 Nfkbie Bcl2 Cxcl3 Ptprcap Gimap6
Blood neutrophils
d
Neu1 Neu2 Neu3 Neu4 Neu5 Neu6 Neu7 Neu8 Neu9 Neu10
Ifitm1 Lrg1 Lyz2 Fgl2 Cst3 Dennd4a Lst1 Retnlg Wfdc21 Mmp8 Lcn2 R3hdm4 Anxa1 Tspo S100a9 S100a8 Lrg1 Wfdc17 Ifitm1 Wfdc21 Tspo Lcn2 Lrg1 Anxa1 Ifitm2 Csdc4 Stfa2l1 Ly6e Trim30a Ifitm3 Slfn1 Samhd1 Pf4 Ngn Gng11 Cla2a Stmp1 Cd9 Myl6 Gpx4 Plek mt−Co1 Atp8b4 Neat1 Ccnd3 Tmcc1 Jmjd1c Arhgap15 Ipcef1 Xylt1 Elmo1 Pde7b Ebf1 Pax5 Btla Bach2 Atf3 Ikzf3 Ralgps2 Man1a Maml2 Ighd
e
day01_MCAo day01_Sham day07_MCAo day07_Sham day14_MCAo day14_Sham
day01−MCAo day01−Sham day07−MCAo day07−Sham day14−MCAo day14−Sham
Chil3 Atp11a Slc24a3 Pam Per2 Gm36723 Flot2 Ahnak Osox1 Ly6g Cd101 Mrps28 Gngt2 Gm34084 Oscp1 Tgm2 Hsp90ab1 Clec4n Dock4 Bank1 Chrm3 Rps6ka1 Polr2j Wfdc17 Steap4 Rbm38 Ckap4 Chst15 G0s2 Rps6ka5 Hmg20a Ap3m1 Thbs1 Vim Morrbid Tspan2 Mrps28 Gm29282 Peak1 Gm16599 Set Rap1gap2 Cebpb Hdc Litaf Il1rap Cux1 Ifitm1 Wfdc17 Pde7b Sipa1l1 Gm16599

**Supplementary Figure 3. Brain and blood neutrophil transcriptomic profile. (a)** *left*: UMAP of neutrophils subset from brain transcriptomes reveals seven clusters; *middle*: UMAP of the same neutrophils coloured by experimental groups; *right*: Stacked bar plot showing the relative proportions of experimental groups across clusters, numbers above the bars indicate the total number of cells in each cluster. **(b, d)** Heatmap of top 10 DEGs across brain (b) or blood (d) neutrophil clusters. For each brain neutrophil cluster (Neu1 – Neu7) or blood neutrophil cluster (Neu1 – Neu10), DEGs were identified by comparing that cluster with all other neutrophils in the same tissue. Top 10 upregulated genes in each cluster are shown in the heatmap. **(c)** Heatmap of top 10 temporal DEG signatures in MCAo neutrophils from brain. DEGs were identified from pairwise comparisons among time points. Top 10 upregulated and downregulated genes from each pairwise comparison are shown. **(e)** Heatmap of top 10 time point-specific blood neutrophil DEGs in MCAo versus sham mice. Differential expression analysis was performed separately at day 1, day 7, and day 14 by comparing neutrophils from MCAo and sham groups within each time point. Top 10 DEGs from each comparison are shown and grouped by the condition in which they were enriched. Heatmap columns represent genes and rows represent clusters (b, d) or experimental groups (c, e). Colour intensity indicates scaled average expression (z-score). Top 10 DEGs were selected based on $|log2FC| \geq log2(1.5)$, pct $\geq 0.75$ (b, d) or 0.1 (c, e), and Bonferroni-adjusted $p < 0.05$, ranked by log2FC.

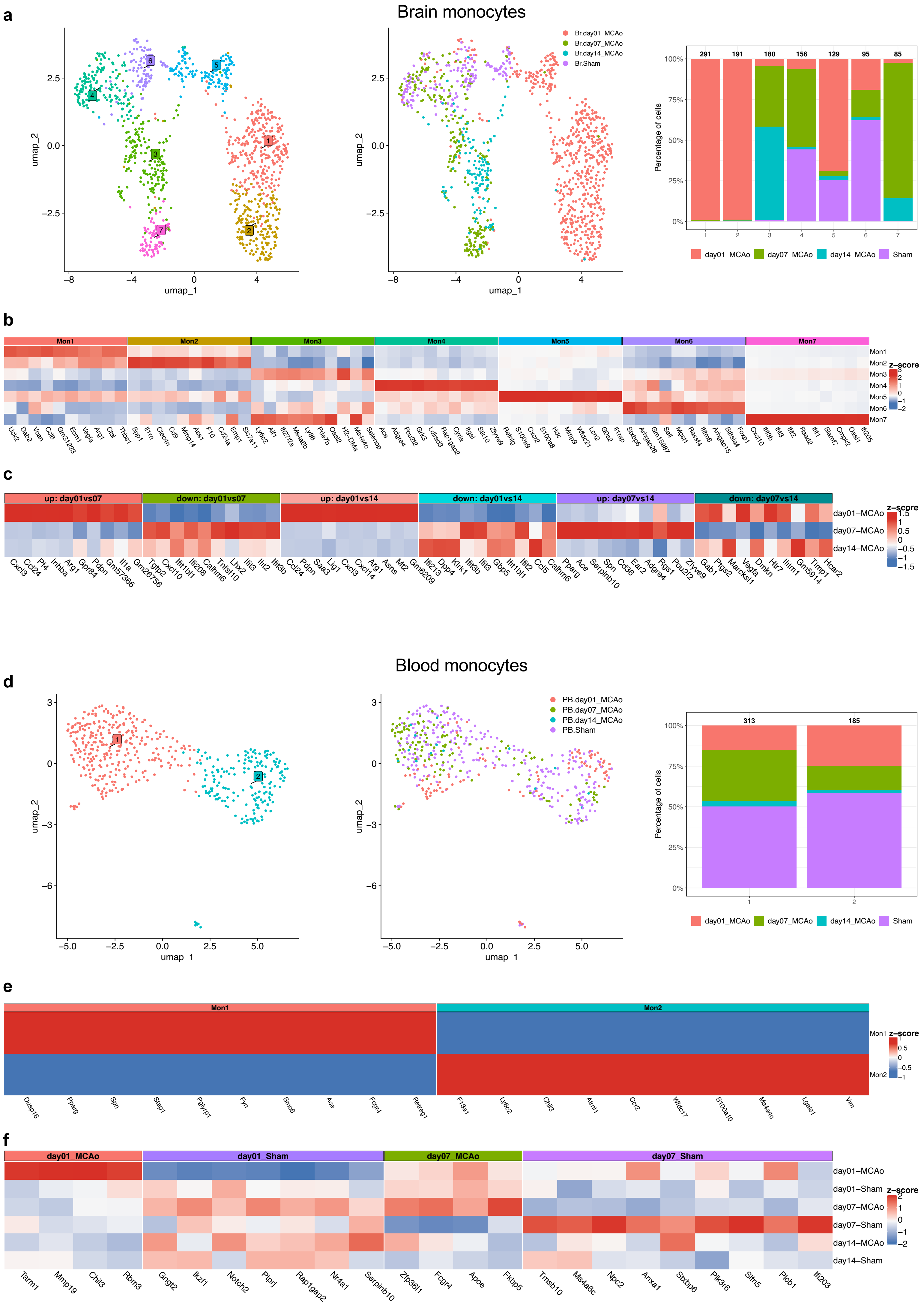
a
Brain monocytes
umap_1
umap_2
Br.day01_MCAo
Br.day07_MCAo
Br.day14_MCAo
Br.Sham
Percentage of cells
291 191 180 156 129 95 85
day01_MCAo
day07_MCAo
day14_MCAo
Sham
b
Mon1 Mon2 Mon3 Mon4 Mon5 Mon6 Mon7
z-score
Uck2 Dab2 Vcan Ccl6 Gm31223 Ecm1 Vegfa Arg1 Ctsl Thbs1 Spp1 Il1rn Clec4n Cd9 Mmp14 Ass1 F10 Cd24a Emp1 Slc7a11 Ly6c2 Aif1 Ifi27l2a Ms4a6b Ly86 Pde7b Oasl2 H2-DMa Ms4a4c Selenop Ace Adgre4 Pou2f2 Grk3 Ldlrad3 Rap1gap2 Cyria Itgal Stk10 Zfyve9 Retnlg S100a9 Cxcr2 S100a8 Hdc Mmp9 Wfdc21 Lcn2 G0s2 Il1rap Stxbp6 Arhgap26 Gm15987 Sell Mgst1 Rassf4 Ifitm6 Arhgap15 St8sia4 Foxp1 Cxcl10 Ifit3b Ifit3 Ifit2 Rsad2 Ifit1 Slamf7 Cmpk2 Oasl1 Ifi205
c
up: day01vs07
down: day01vs07
up: day01vs14
down: day01vs14
up: day07vs14
down: day07vs14
day01−MCAo
day07−MCAo
day14−MCAo
z-score
Cxcl3 Ccl24 Pf4 Inhba Arg1 Gpr84 Pdpn Gm57366 Il1a Gm26756 Tgtp2 Cxcl10 Ifit1bl1 Ifi208 Calhm6 Tnfsf10 Lhx2 Ifit3 Ifit2 Ifit3b Ccl24 Pdpn Saa3 Lig1 Cxcl3 Cxcl14 Arg1 Asns Mt2 Gm6209 Ifi213 Dpp4 Klrk1 Ifit3b Ifit3 Gbp5 Ifit1bl1 Ifit2 Ccl5 Calhm6 Pparg Ace Serpinb10 Spn Cd36 Ear2 Adgre4 Rgs1 Pou2f2 Zfyve9 Gab1 Ptgs2 Marcksl1 Vegfa Dmkn Htr7 Ifitm1 Gm5914 Timp1 Hcar2
Blood monocytes
d
umap_1
umap_2
PB.day01_MCAo
PB.day07_MCAo
PB.day14_MCAo
PB.Sham
Percentage of cells
313 185
day01_MCAo
day07_MCAo
day14_MCAo
Sham
e
Mon1 Mon2
z-score
Dusp16 Pparg Spn Stap1 Pglyrp1 Fyn Smc6 Ace Fcgr4 Retreg1 F13a1 Ly6c2 Chil3 Atrnl1 Ccr2 Wfdc17 S100a10 Ms4a4c Lgals1 Vim
f
day01_MCAo
day01_Sham
day07_MCAo
day07_Sham
day01−MCAo
day01−Sham
day07−MCAo
day07−Sham
day14−MCAo
day14−Sham
z-score
Tarm1 Mmp19 Chil3 Rbm3 Gngt2 Ikzf1 Notch2 Ptprj Rap1gap2 Nr4a1 Serpinb10 Zfp36l1 Fcgr4 Apoe Fkbp5 Tmsb10 Ms4a6c Npc2 Anxa1 Stxbp6 Pik3r6 Slfn5 Plcb1 Ifi203

**Supplementary Figure 4. Brain and blood monocyte transcriptomic profile. (a, d)** *left*: UMAP of monocytes subset from brain (a) or blood (d) transcriptomes reveals seven (brain) or two (blood) clusters; *middle*: UMAP of the same monocytes coloured by experimental groups; *right*: Stacked bar plot showing the relative proportions of experimental groups across clusters, numbers above the bars indicate the total number of cells in each cluster. **(b, e)** Heatmap of top 10 DEGs across brain (b) or blood (e) monocyte clusters. For each brain monocyte cluster (Mon1 – Mon7) or blood monocyte cluster (Mon1 – Mon2), DEGs were identified by comparing that cluster with all other monocytes in the same tissue. Top 10 upregulated genes in each cluster are shown in the heatmap. **(c)** Heatmap of top 10 temporal DEG signatures in MCAo monocytes from brain. DEGs were identified from pairwise comparisons among time points. Top 10 upregulated and downregulated genes from each pairwise comparison are shown. **(f)** Heatmap of top 10 time point-specific blood monocytes DEGs in MCAo versus sham mice. Differential expression analysis was performed separately at day 1, day 7, and day 14 by comparing monocytes from MCAo and sham groups within each time point. Top 10 DEGs from each comparison are shown and grouped by the condition in which they were enriched. Heatmap columns represent genes and rows represent clusters (b, e) or experimental groups (c, f). Colour intensity indicates scaled average expression (z-score). Top 10 DEGs were selected based on $|\log2FC| \geq \log2(1.5)$, pct $\geq 0.75$ (b, e) or 0.1 (c, f), and Bonferroni-adjusted $p < 0.05$, ranked by log2FC.

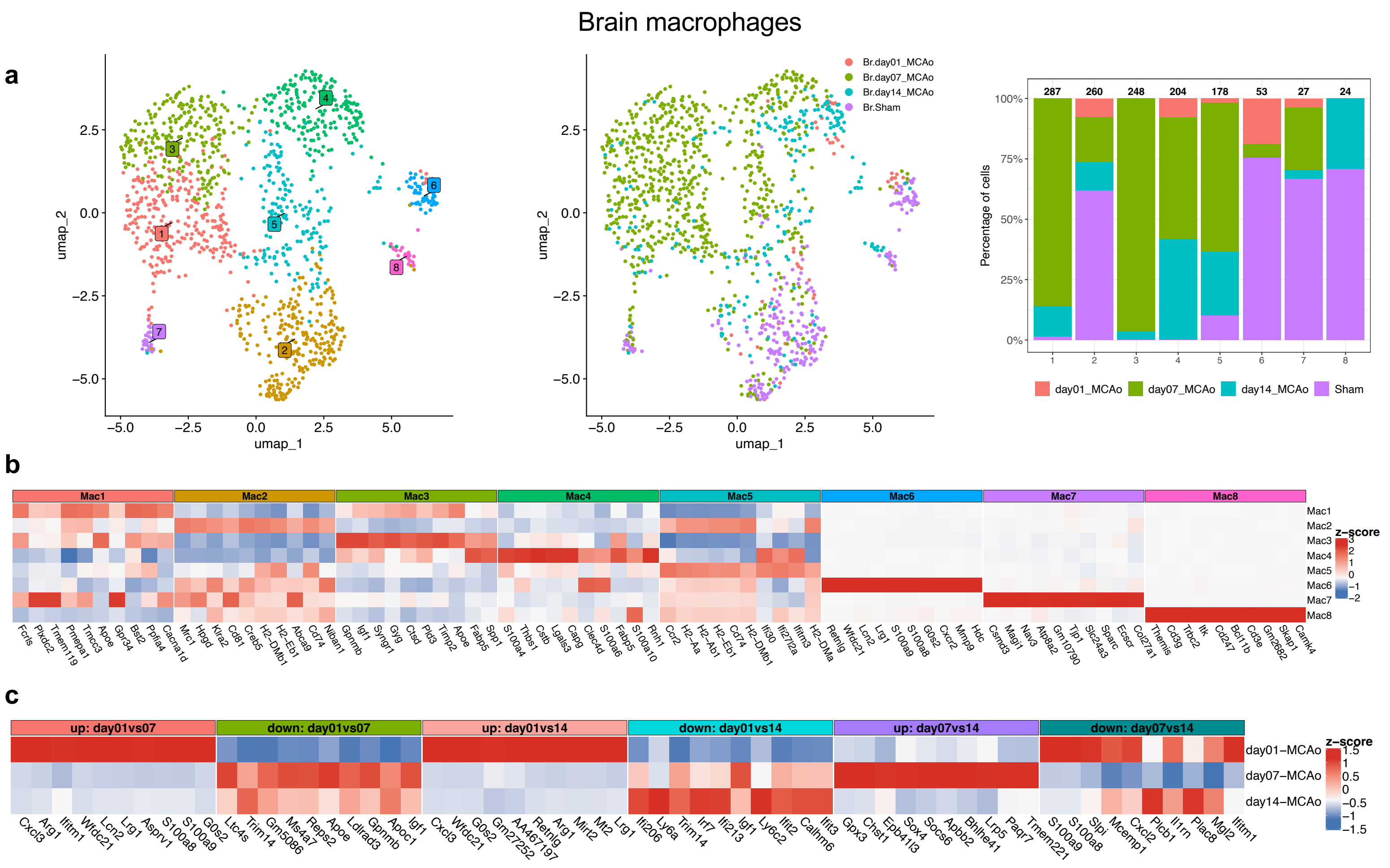
Brain macrophages
a
b
c
umap_1
umap_2
Br.day01_MCAo
Br.day07_MCAo
Br.day14_MCAo
Br.Sham
Percentage of cells
287 260 248 204 178 53 27 24
day01_MCAo day07_MCAo day14_MCAo Sham
Mac1 Mac2 Mac3 Mac4 Mac5 Mac6 Mac7 Mac8
z-score
up: day01vs07
down: day01vs07
up: day01vs14
down: day01vs14
up: day07vs14
down: day07vs14
day01-MCAo
day07-MCAo
day14-MCAo

**Supplementary Figure 5. Brain macrophage transcriptomic profile. (a)** *left*: UMAP of macrophages subset from brain transcriptomes reveals eight clusters; *middle*: UMAP of the same macrophages coloured by experimental groups; *right*: Stacked bar plot showing the relative proportions of experimental groups across clusters, numbers above the bars indicate the total number of cells in each cluster. **(b)** Heatmap of top 10 DEGs across brain macrophage clusters. For each brain macrophage cluster (Mac1 – Mac8), DEGs were identified by comparing that cluster with all other brain macrophages. Top 10 upregulated genes in each cluster are shown in the heatmap. **(c)** Heatmap of top 10 temporal DEG signatures in MCAo macrophages from brain. DEGs were identified from pairwise comparisons among time points. Top 10 upregulated and downregulated genes from each pairwise comparison are shown. Heatmap columns represent genes and rows represent clusters (b) or experimental groups (c). Colour intensity indicates scaled average expression (z-score). Top 10 DEGs were selected based on $|log2FC| \geq log2(1.5)$, pct $\geq 0.75$ (b) or 0.1 (c), and Bonferroni-adjusted $p < 0.05$, ranked by log2FC.

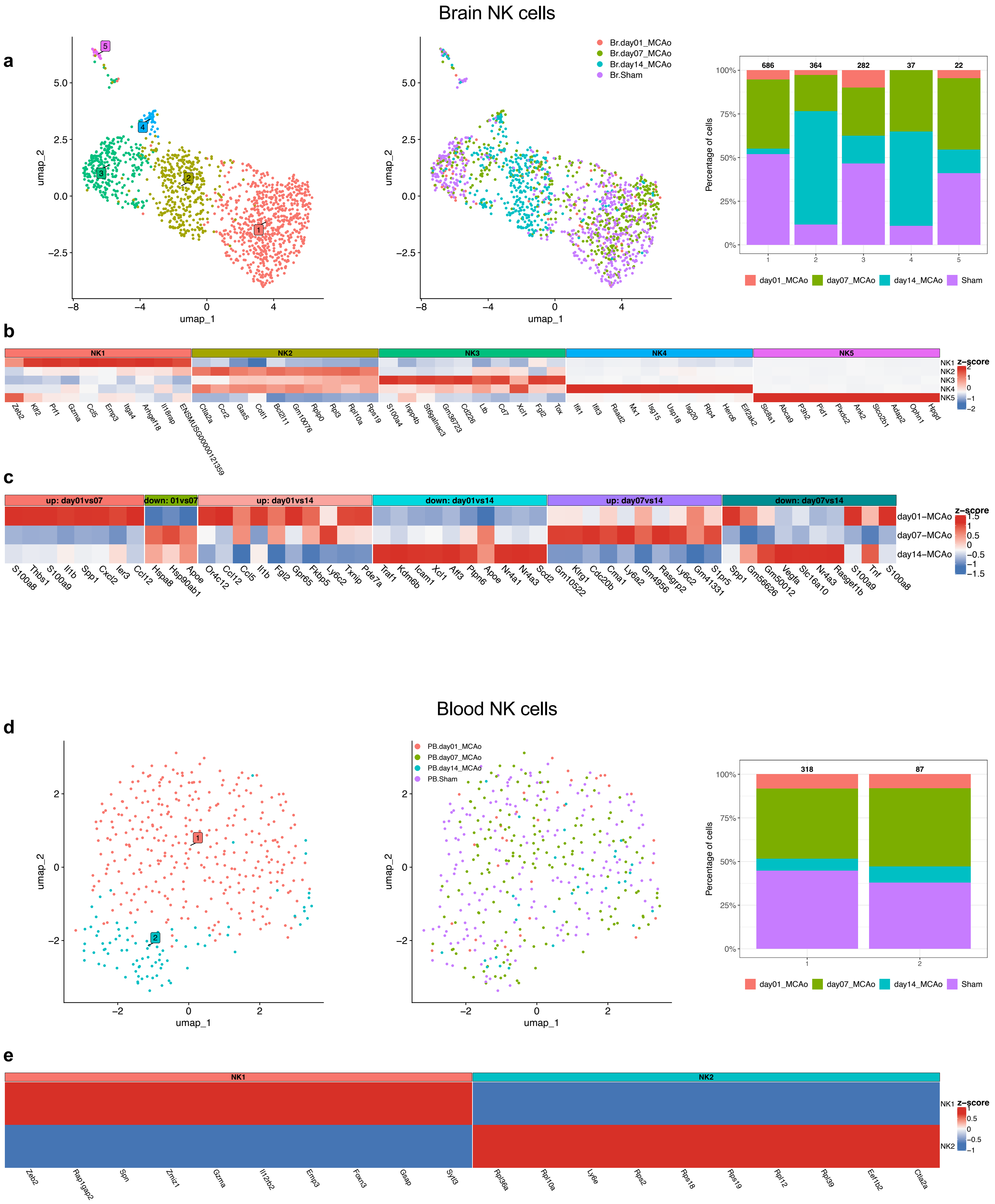

Brain NK cells
a
umap_1
umap_2
Br.day01_MCAo
Br.day07_MCAo
Br.day14_MCAo
Br.Sham
Percentage of cells
686
364
282
37
22
day01_MCAo
day07_MCAo
day14_MCAo
Sham
b
NK1
NK2
NK3
NK4
NK5
z-score
Zeb2 Klf2 Prf1 Gzma Ccl5 Emp3 Itga4 Arhgef18 Il18rap ENSMUSG00000121359
Ctla2a Ccr2 Gas5 Cd7 Bcl2l11 Gm10076 Rplp0 Rpl3 Rpl10a Rps19
S100a4 Inpp4b St6galnac3 Gm36723 Cd226 Ltb Ccl7 Xcl1 Fgl2 Tox
Ifit1 Ifit3 Rsad2 Mx1 Isg15 Usp18 Isg20 Rtp4 Herc6 Eif2ak2
Slc8a1 Abca9 P3h2 Pld1 Pkdc2 Ank2 Slco2b1 Adap2 Ophn1 Hpgd
c
up: day01vs07
down: 01vs07
up: day01vs14
down: day01vs14
up: day07vs14
down: day07vs14
day01−MCAo
day07−MCAo
day14−MCAo
z-score
S100a8 Thbs1 S100a9 Il1b Spp1 Cxcl2 Ier3 Ccl12 Hspa8 Hsp90ab1 Apoe Or4c12 Ccl12 Ccl5 Il1b Fgl2 Gpr65 Fkbp5 Ly6c2 Txnip Pde7a Traf1 Kdm6b Icam1 Xcl1 Aff3 Ptpn6 Apoe Nr4a1 Nr4a3 Scd2 Gm10522 Klrg1 Cdc20b Cma1 Ly6a2 Gm4956 Rasgrp2 Ly6c2 Gm41331 S1pr5 Spp1 Gm56626 Gm50012 Vegfa Slc16a10 Nr4a3 Rasgef1b S100a9 Tnf S100a8
Blood NK cells
d
umap_1
umap_2
PB.day01_MCAo
PB.day07_MCAo
PB.day14_MCAo
PB.Sham
Percentage of cells
318
87
day01_MCAo
day07_MCAo
day14_MCAo
Sham
e
NK1
NK2
z-score
Zeb2 Rap1gap2 Spn Zmiz1 Gzma Il12rb2 Emp3 Foxn3 Gsap Sytl3 Rpl36a Rpl10a Ly6e Rps2 Rps18 Rps19 Rpl12 Rpl39 Eef1b2 Ctla2a

**Supplementary Figure 6. Brain and blood natural killer (NK) cell transcriptomic profile. (a, d)** *left*: UMAP of NK cells subset from brain (a) or blood (d) transcriptomes reveals five (brain) or two (blood) clusters; *middle*: UMAP of the same NK cells coloured by experimental groups; *right*: Stacked bar plot showing the relative proportions of experimental groups across clusters, numbers above the bars indicate the total number of cells in each cluster. **(b, e)** Heatmap of top 10 DEGs across brain (b) or blood (e) NK cell clusters. For each brain NK cell cluster (NK1 – NK5) or blood NK cell cluster (NK1 – NK2), DEGs were identified by comparing that cluster with all other NK cells in the same tissue. Top 10 upregulated genes in each cluster are shown in the heatmap. **(c)** Heatmap of top 10 temporal DEG signatures in MCAo NK cells from brain. DEGs were identified from pairwise comparisons among time points. Top 10 upregulated and downregulated genes from each pairwise comparison are shown. Heatmap columns represent genes and rows represent clusters (b, e) or experimental groups (c). Colour intensity indicates scaled average expression (z-score). Top 10 DEGs were selected based on $|\log2FC| \geq \log2(1.5)$, pct $\geq 0.75$ (b, e) or 0.1 (c), and Bonferroni-adjusted $p < 0.05$, ranked by log2FC.

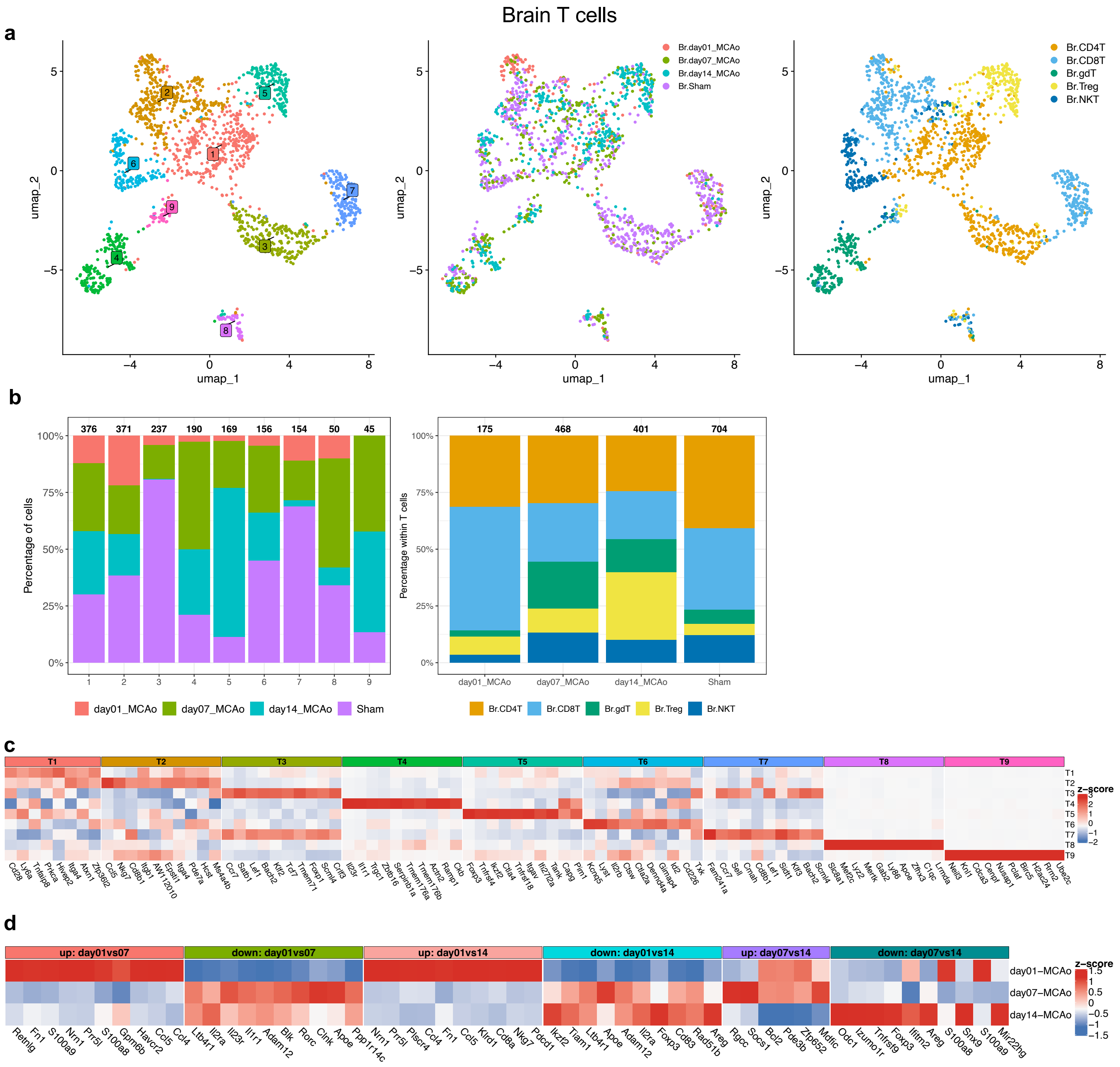
Brain T cells
a
umap_2
umap_1
Br.day01_MCAo
Br.day07_MCAo
Br.day14_MCAo
Br.Sham
Br.CD4T
Br.CD8T
Br.gdT
Br.Treg
Br.NKT
b
Percentage of cells
376 371 237 190 169 156 154 50 45
day01_MCAo day07_MCAo day14_MCAo Sham
Percentage within T cells
175 468 401 704
day01_MCAo day07_MCAo day14_MCAo Sham
Br.CD4T Br.CD8T Br.gdT Br.Treg Br.NKT
c
T1 T2 T3 T4 T5 T6 T7 T8 T9
z-score
d
up: day01vs07
down: day01vs07
up: day01vs14
down: day01vs14
up: day07vs14
down: day07vs14
day01−MCAo
day07−MCAo
day14−MCAo
z-score

**Supplementary Figure 7. Brain T cell transcriptomic profile. (a)** *left*: UMAP of T cells subset from brain transcriptomes reveals nine clusters; *middle*: UMAP of the same T cells coloured by experimental groups; *right*: UMAP of the same T cells coloured by T cell subpopulations. **(b)** *left*: Stacked bar plot showing the relative proportions of experimental groups across clusters, numbers above the bars indicate the total number of cells in each cluster; *right*: Stacked bar plot showing the relative proportions of T cell subpopulations across experimental groups, numbers above the bars indicate the total number of cells in each group. **(c)** Heatmap of top 10 DEGs across brain T cell clusters. For each brain T cell cluster (T1 – T9), DEGs were identified by comparing that cluster with all other brain T cells. Top 10 upregulated genes in each cluster are shown in the heatmap. **(d)** Heatmap of top 10 temporal DEG signatures in MCAo T cells from brain. DEGs were identified from pairwise comparisons among time points. Top 10 upregulated and downregulated genes from each pairwise comparison are shown. Heatmap columns represent genes and rows represent clusters (c) or experimental groups (d). Colour intensity indicates scaled average expression (z-score). Top 10 DEGs were selected based on $|log2FC| \geq log2(1.5)$, pct ≥ 0.75 (c) or 0.1 (d), and Bonferroni-adjusted $p < 0.05$, ranked by log2FC.

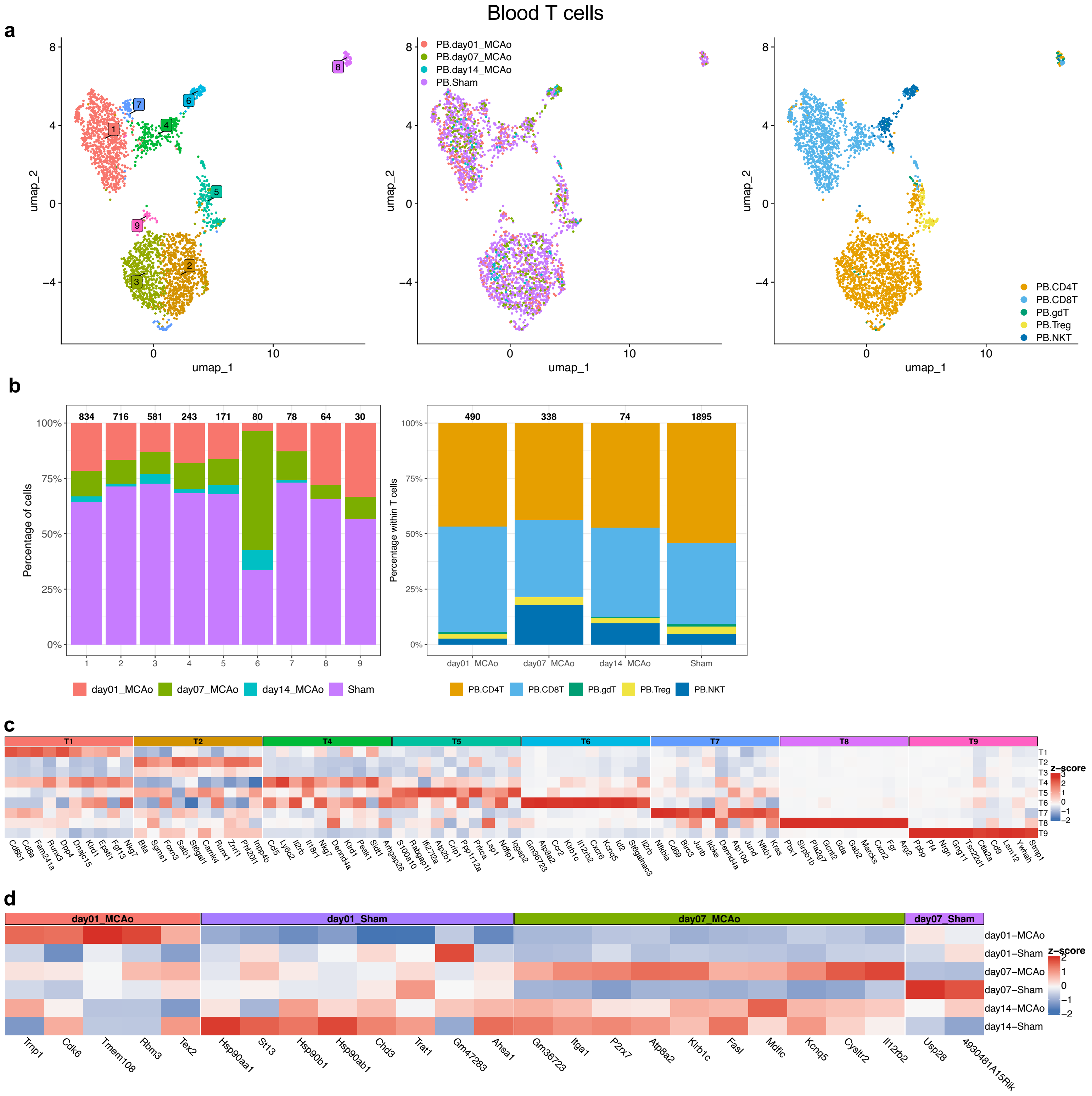

Blood T cells
a
umap_1
umap_2
PB.day01_MCAo
PB.day07_MCAo
PB.day14_MCAo
PB.Sham
PB.CD4T
PB.CD8T
PB.gdT
PB.Treg
PB.NKT
b
834 716 581 243 171 80 78 64 30
Percentage of cells
day01_MCAo day07_MCAo day14_MCAo Sham
490 338 74 1895
Percentage within T cells
day01_MCAo day07_MCAo day14_MCAo Sham
c
T1 T2 T4 T5 T6 T7 T8 T9
z-score
d
day01_MCAo day01_Sham day07_MCAo day07_Sham
day01–MCAo
day01–Sham
day07–MCAo
day07–Sham
day14–MCAo
day14–Sham
z-score
Trnp1 Cdk6 Tmem108 Rbm3 Tex2 Hsp90aa1 St13 Hsp90b1 Hsp90ab1 Chd3 Trat1 Gm47283 Ahsa1 Gm36723 Itga1 P2rx7 Atp8a2 Klrb1c Fasl Mdfic Kcnq5 Cysltr2 Il12rb2 Usp28 4930481A15Rik

**Supplementary Figure 8. Blood T cell transcriptomic profile. (a)** *left*: UMAP of T cells subset from blood transcriptomes reveals nine clusters; *middle*: UMAP of the same T cells coloured by experimental groups; *right*: UMAP of the same T cells coloured by T cell subpopulations. **(b)** *left*: Stacked bar plot showing the relative proportions of experimental groups across clusters, numbers above the bars indicate the total number of cells in each cluster; *right*: Stacked bar plot showing the relative proportions of T cell subpopulations across experimental groups, numbers above the bars indicate the total number of cells in each group. **(c)** Heatmap of top 10 DEGs across blood T cell clusters. For each blood T cell cluster (T1 – T9), DEGs were identified by comparing that cluster with all other blood T cells. Top 10 upregulated genes in each cluster are shown in the heatmap. **(d)** Heatmap of top 10 time point-specific blood T cell DEGs in MCAo versus sham mice. Differential expression analysis was performed separately at day 1, day 7, and day 14 by comparing T cells from MCAo and sham groups within each time point. Top 10 DEGs from each comparison are shown and grouped by the condition in which they were enriched. Heatmap columns represent genes and rows represent clusters (c) or experimental groups (d). Colour intensity indicates scaled average expression (z-score). Top 10 DEGs were selected based on $|log2FC| \geq log2(1.5)$, pct ≥ 0.75 (c) or 0.1 (d), and Bonferroni-adjusted $p < 0.05$, ranked by log2FC.

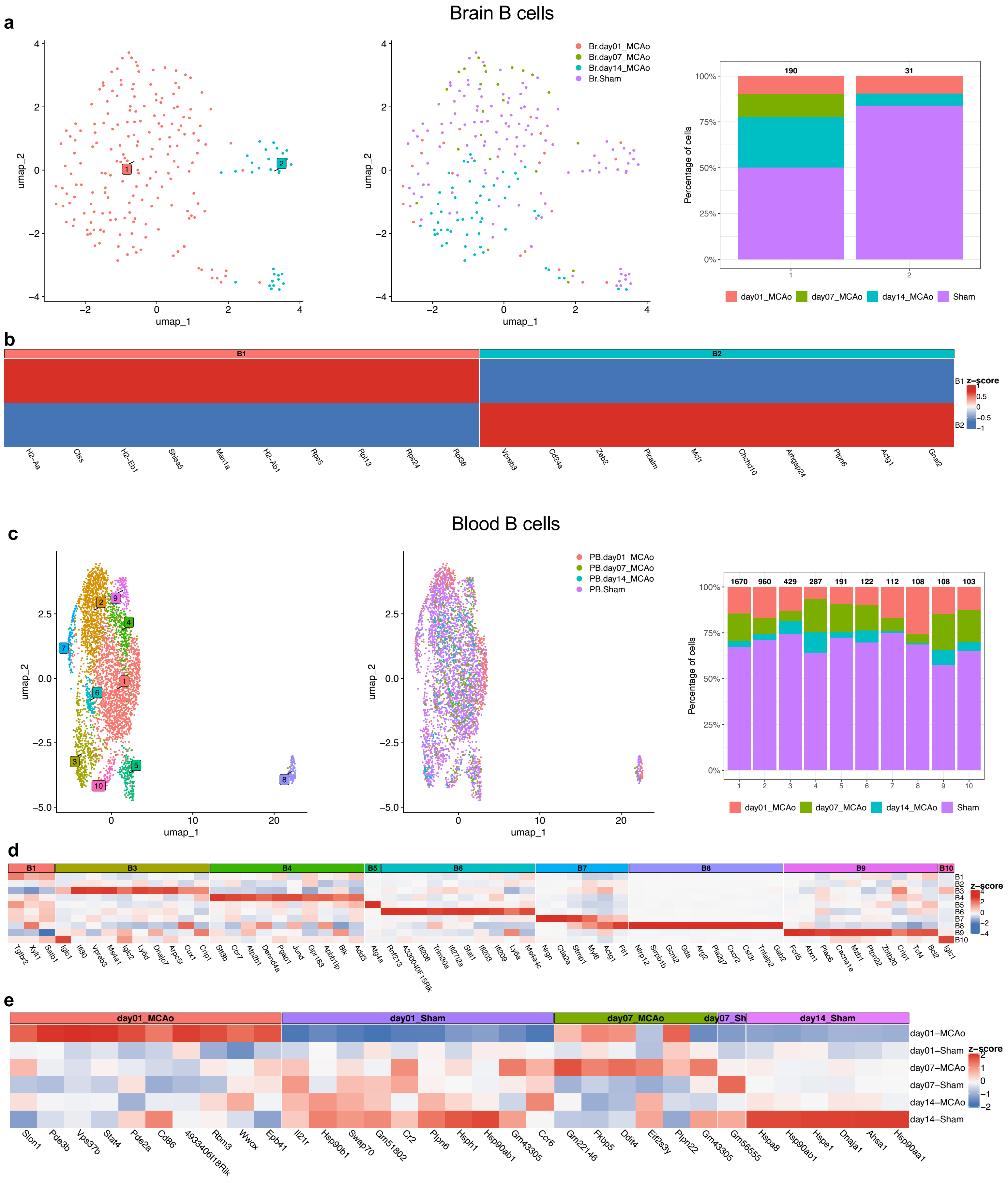

a
Brain B cells
Br.day01_MCAo
Br.day07_MCAo
Br.day14_MCAo
Br.Sham
umap_1
umap_2
Percentage of cells
190
31
day01_MCAo
day07_MCAo
day14_MCAo
Sham
b
B1
B2
z-score
H2-Aa
Ctss
H2-Eb1
Shisa5
Man1a
H2-Ab1
Rps5
Rpl13
Rps24
Rpl36
Vpreb3
Cd24a
Zeb2
Picalm
Mcl1
Chchd10
Arhgap24
Ptpn6
Actg1
Gnai2
c
Blood B cells
PB.day01_MCAo
PB.day07_MCAo
PB.day14_MCAo
PB.Sham
1670
960
429
287
191
122
112
108
108
103
d
B1
B3
B4
B5
B6
B7
B8
B9
B10
e
day01_MCAo
day01_Sham
day07_MCAo
day07_Sh
day14_Sham
day01–MCAo
day01–Sham
day07–MCAo
day07–Sham
day14–MCAo
day14–Sham
Ston1
Pde3b
Vps37b
Stat4
Pde2a
Cd86
4933406I18Rik
Rbm3
Wwox
Epb41
Il21r
Hsp90b1
Swap70
Gm51802
Cr2
Ptpn6
Hsph1
Hsp90ab1
Gm43305
Ccr6
Gm22146
Fkbp5
Ddit4
Eif2s3y
Ptpn22
Gm43305
Gm56555
Hspa8
Hsp90ab1
Hspe1
Dnaja1
Ahsa1
Hsp90aa1

**Supplementary Figure 9. Brain and blood B cell transcriptomic profile. (a, c)** *left*: UMAP of B cells subset from brain (a) or blood (c) transcriptomes reveals two (brain) or ten (blood) clusters; *middle*: UMAP of the same B cells coloured by experimental groups; *right*: Stacked bar plot showing the relative proportions of experimental groups across clusters, numbers above the bars indicate the total number of cells in each cluster. **(b, d)** Heatmap of top 10 DEGs across brain (b) or blood (e) B cell clusters. For each brain B cell cluster (B1 – B2) or blood B cell cluster (B1 – B10), DEGs were identified by comparing that cluster with all other B cells in the same tissue. Top 10 upregulated genes in each cluster are shown in the heatmap. **(e)** Heatmap of top 10 time point-specific blood B cell DEGs in MCAo versus sham mice. Differential expression analysis was performed separately at day 1, day 7, and day 14 by comparing B cells from MCAo and sham groups within each time point. Top 10 DEGs from each comparison are shown and grouped by the condition in which they were enriched. Heatmap columns represent genes and rows represent clusters (b, d) or experimental groups (e). Colour intensity indicates scaled average expression (z-score). Top 10 DEGs were selected based on $|log2FC| \geq log2(1.5)$, $pct \geq 0.75$ (b, d) or 0.1 (e), and Bonferroni-adjusted $p < 0.05$, ranked by log2FC.

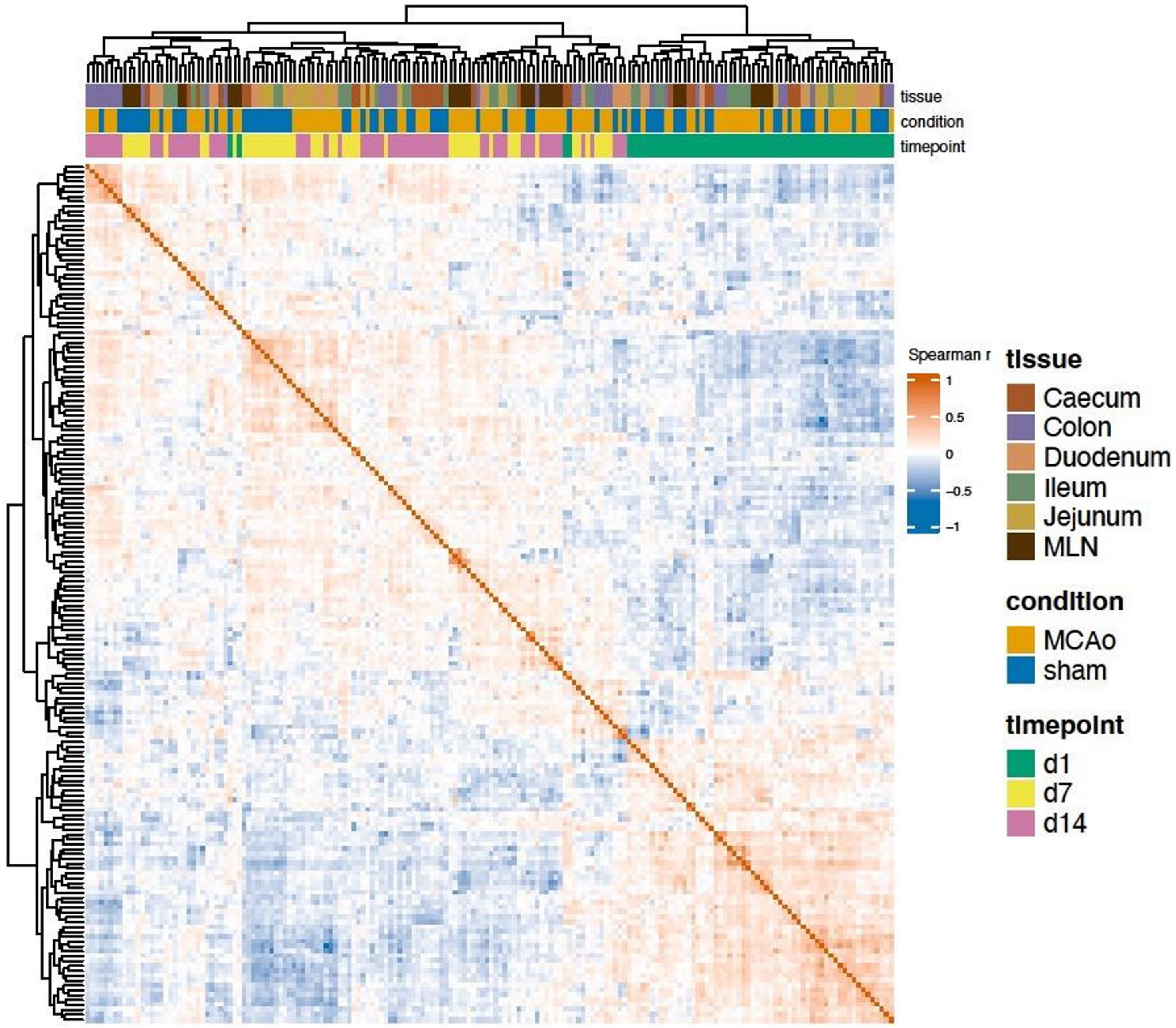
tissue
condition
timepoint
Spearman r
1
0.5
0
−0.5
−1
tissue
Caecum
Colon
Duodenum
Ileum
Jejunum
MLN
condition
MCAo
sham
timepoint
d1
d7
d14

**Supplementary Figure 10. Sample-to-sample correlation heatmap of intestinal proteomes after SVA batch correction.** Spearman correlation matrix across all 176 samples (6 tissues × 3 timepoints × MCAo/sham), computed on SVA-corrected intensities z-scored per tissue and restricted to the 4,079 proteins detected in all six tissues. Rows and columns are hierarchically clustered (Euclidean distance, complete linkage). Top annotations indicate tissue (caecum, colon, duodenum, ileum, jejunum, MLN), condition (MCAo, sham), and timepoint (d1, d7, d14).

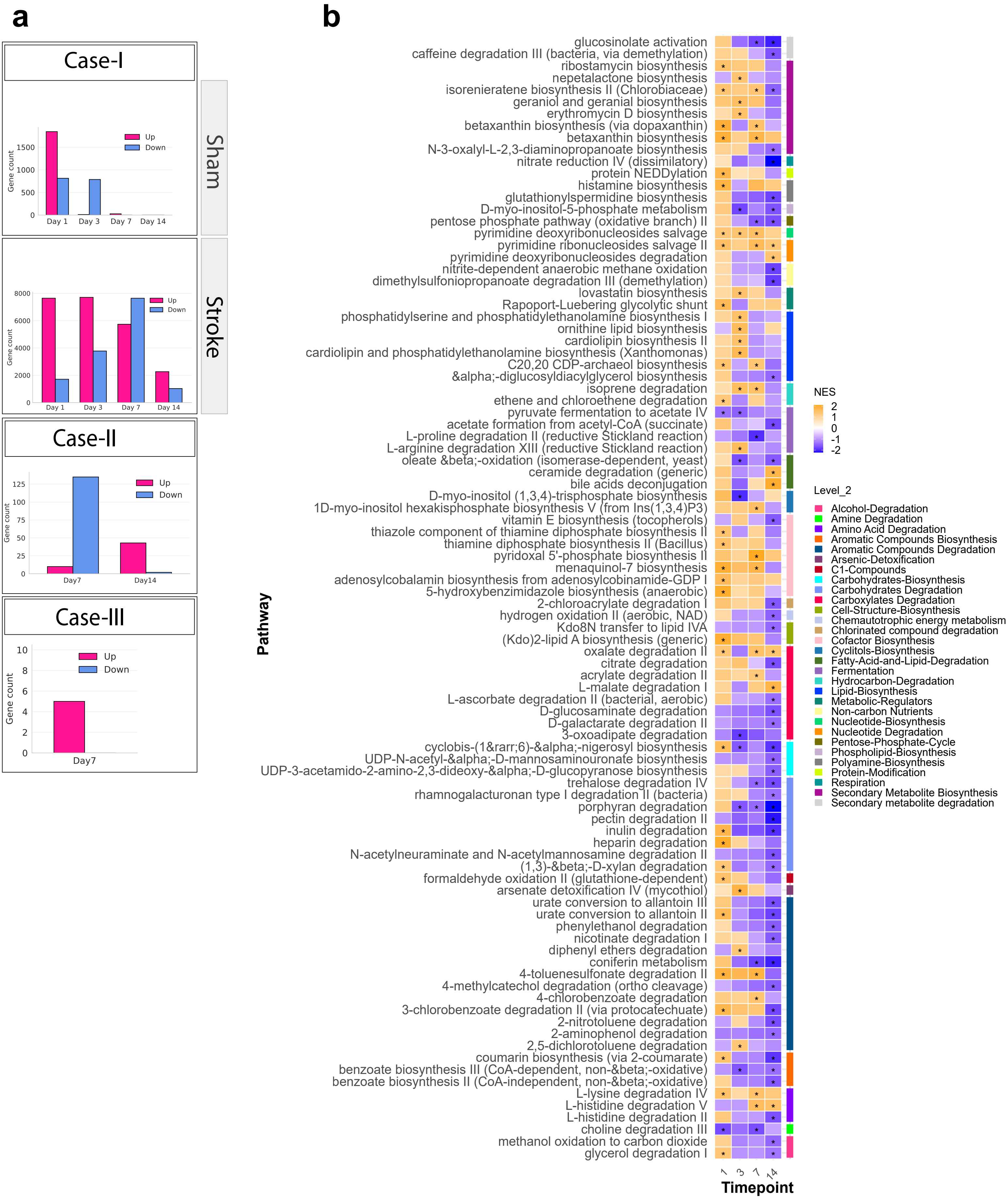

a
Case-I
Sham
Stroke
Gene count
Up
Down
Day 1
Day 3
Day 7
Day 14
Case-II
Day7
Day14
Case-III
b
glucosinolate activation
caffeine degradation III (bacteria, via demethylation)
ribostamycin biosynthesis
nepetalactone biosynthesis
isorenieratene biosynthesis II (Chlorobiaceae)
geraniol and geranial biosynthesis
erythromycin D biosynthesis
betaxanthin biosynthesis (via dopaxanthin)
betaxanthin biosynthesis
N-3-oxalyl-L-2,3-diaminopropanoate biosynthesis
nitrate reduction IV (dissimilatory)
protein NEDDylation
histamine biosynthesis
glutathionylspermidine biosynthesis
D-myo-inositol-5-phosphate metabolism
pentose phosphate pathway (oxidative branch) II
pyrimidine deoxyribonucleosides salvage
pyrimidine ribonucleosides salvage II
pyrimidine deoxyribonucleosides degradation
nitrite-dependent anaerobic methane oxidation
dimethylsulfoniopropanoate degradation III (demethylation)
lovastatin biosynthesis
Rapoport-Luebering glycolytic shunt
phosphatidylserine and phosphatidylethanolamine biosynthesis I
ornithine lipid biosynthesis
cardiolipin biosynthesis II
cardiolipin and phosphatidylethanolamine biosynthesis (Xanthomonas)
C20,20 CDP-archaeol biosynthesis
α-diglucosyldiacylglycerol biosynthesis
isoprene degradation
ethene and chloroethene degradation
pyruvate fermentation to acetate IV
acetate formation from acetyl-CoA (succinate)
L-proline degradation II (reductive Stickland reaction)
L-arginine degradation XIII (reductive Stickland reaction)
oleate β-oxidation (isomerase-dependent, yeast)
ceramide degradation (generic)
bile acids deconjugation
D-myo-inositol (1,3,4)-trisphosphate biosynthesis
1D-myo-inositol hexakisphosphate biosynthesis V (from Ins(1,3,4)P3)
vitamin E biosynthesis (tocopherols)
thiazole component of thiamine diphosphate biosynthesis II
thiamine diphosphate biosynthesis II (Bacillus)
pyridoxal 5'-phosphate biosynthesis II
menaquinol-7 biosynthesis
adenosylcobalamin biosynthesis from adenosylcobinamide-GDP I
5-hydroxybenzimidazole biosynthesis (anaerobic)
2-chloroacrylate degradation I
hydrogen oxidation II (aerobic, NAD)
Kdo8N transfer to lipid IVA
(Kdo)2-lipid A biosynthesis (generic)
oxalate degradation II
citrate degradation
acrylate degradation II
L-malate degradation I
L-ascorbate degradation II (bacterial, aerobic)
D-glucosaminate degradation
D-galactarate degradation II
3-oxoadipate degradation
cyclobis-(1→6)-α-nigerosyl biosynthesis
UDP-N-acetyl-α-D-mannosaminouronate biosynthesis
UDP-3-acetamido-2-amino-2,3-dideoxy-α-D-glucopyranose biosynthesis
trehalose degradation IV
rhamnogalacturonan type I degradation II (bacteria)
porphyran degradation
pectin degradation II
inulin degradation
heparin degradation
N-acetylneuraminate and N-acetylmannosamine degradation II
(1,3)-β-D-xylan degradation
formaldehyde oxidation II (glutathione-dependent)
arsenate detoxification IV (mycothiol)
urate conversion to allantoin III
urate conversion to allantoin II
phenylethanol degradation
nicotinate degradation I
diphenyl ethers degradation
coniferin metabolism
4-toluenesulfonate degradation II
4-methylcatechol degradation (ortho cleavage)
4-chlorobenzoate degradation
3-chlorobenzoate degradation II (via protocatechuate)
2-nitrotoluene degradation
2-aminophenol degradation
2,5-dichlorotoluene degradation
coumarin biosynthesis (via 2-coumarate)
benzoate biosynthesis III (CoA-dependent, non-β-oxidative)
benzoate biosynthesis II (CoA-independent, non-β-oxidative)
L-lysine degradation IV
L-histidine degradation V
L-histidine degradation II
choline degradation III
methanol oxidation to carbon dioxide
glycerol degradation I
Pathway
Timepoint
NES
Level_2
Alcohol-Degradation
Amine Degradation
Amino Acid Degradation
Aromatic Compounds Biosynthesis
Aromatic Compounds Degradation
Arsenic-Detoxification
C1-Compounds
Carbohydrates-Biosynthesis
Carbohydrates Degradation
Carboxylates Degradation
Cell-Structure-Biosynthesis
Chemautotrophic energy metabolism
Chlorinated compound degradation
Cofactor Biosynthesis
Cyclitols-Biosynthesis
Fatty-Acid-and-Lipid-Degradation
Fermentation
Hydrocarbon-Degradation
Lipid-Biosynthesis
Metabolic-Regulators
Non-carbon Nutrients
Nucleotide-Biosynthesis
Nucleotide Degradation
Pentose-Phosphate-Cycle
Phospholipid-Biosynthesis
Polyamine-Biosynthesis
Protein-Modification
Respiration
Secondary Metabolite Biosynthesis
Secondary metabolite degradation

**Supplementary Figure 11. Differential expression analysis and enrichment analysis. (a)** Three approaches using dream (differential expression for repeated measures) to study differential expression analysis. Case-I (within-condition temporal contrasts) comparing each timepoint to day 0 (baseline) within stroke and sham separately, Case-II (cross-sectional contrasts) comparing stroke to sham at each individual timepoint, and Case-III (interaction contrasts) formally testing whether the change from baseline differed between stroke and sham at each timepoint. **(b)** Gut microbiome transcriptional enrichment of MetaCyc pathways across timepoints following stroke in Case-II. Heatmap displays normalized enrichment scores (NES) from gene set enrichment analysis (GSEA) of non-redundant MetaCyc pathways at days 1, 3, 7, and 14 following stroke. Color intensity reflects NES magnitude with warm colors (orange) indicate stroke-enriched pathways (NES > 0 & padj < 0.05) and cool colors (purple) indicate sham-enriched pathways (NES < 0 & padj < 0.05). Asterisks denote pathways passing the significance threshold. Pathway redundancy was minimized using the collapse function from the *fgsea* package. Right-side color bar indicates MetaCyc level-2 functional hierarchy classification.
